# A synthetic xeno-free extracellular matrix surrogate supports robust human induced pluripotent stem cell maintenance and multilineage differentiation

**DOI:** 10.64898/2026.10.07.757018

**Authors:** Constantin Berger, Linus Wiora, Terri L Holmes, Fabiola Walz, Clemens Sauter, Luisa Baraban, Julian Schwebler, Geraldine Beer, Sonja Lauke, Jana Keller, Melanie Kraft, Yvonne Schelling, Ka Cheung, Elina Nürnberg, Sarah Al Muthafer, Sandra Segeletz, Christina Neser, James G.W. Smith, Stefan Hauser, Daniela Zdzieblo

**Affiliations:** Translational Center Regenerative Therapies TLC-RT, Fraunhofer Institute for Silicate Research ISC, Würzburg, Bavaria, Germany; German Center for Neurodegenerative Diseases (DZNE), Tübingen, Baden-Württemberg, Germany; Department of Neurology and Hertie Institute for Clinical Brain Research, University of Tübingen, Tübingen, Baden-Württemberg, Germany; Graduate School of Cellular and Molecular Neuroscience, University of Tübingen, Tübingen, Baden-Württemberg, Germany; Norwich Medical School, University of East Anglia, Norwich, Norfolk, UK; Department for Functional Materials in Medicine and Dentistry, University Hospital Würzburg, Würzburg, Bavaria, Germany; Department of General, Visceral, Transplantation, Vascular and Paediatric Surgery (Department of Surgery I), University Hospital Würzburg, Würzburg, Bavaria, Germany; faCellitate GmbH, Mannheim, Baden-Württemberg, Germany; Gene and RNA Therapy Center (GRTC), Faculty of Medicine, University of Tübingen, Tübingen, Baden-Württemberg, Germany

**Keywords:** Stem Cell culture, hiPSCs, ECM coating, synthetic coating

## Abstract

Animal-derived surface coatings present the gold standard in stem cell culture. However, they come with major drawbacks such as lot-to-lot inconsistency and an ethically problematic production. Approaches to develop synthetic coating alternatives have largely failed to recapitulate biological functions essential to PSC-self renewal and differentiation. In this study, we have investigated the suitability of a fully synthetic, chemically defined extracellular matrix-mimicking coating named BIOINSTRUCT^TM^ for hiPSC culture. BIOINSTRUCT^TM^ was employed during maintenance culture of IMR90-4 cells, as well as directed differentiation of HIHCNi008-A, ACTN2-GFP, and UKKI-011-A cells. BIOINSTRUCT^TM^ preserved hiPSC characteristics over the experimental period of 10 passages similar to Matrigel^TM^. Spontaneous differentiation resulted in the formation of ectodermal, mesodermal and endodermal derivatives, confirming the retention of pluripotency. Directed differentiation on BIOINSTRUCT^TM^ enabled the generation of neuronal, cardiac and pancreatic endocrine cells. Altogether, our data underscore BIOINSTRUCT^TM^ as a promising alternative to animal-derived extracellular matrix coatings.

## Introduction

Human induced pluripotent stem cells (hiPSCs) have become indispensable tools in modern biomedical research due to their capacity for unlimited self-renewal and differentiation into virtually all somatic cell types. Since their initial description^1,2^, hiPSCs have enabled transformative advances in regenerative medicine, offering prospects for patient-specific cell replacement therapies in disorders such as Parkinson’s disease^3^, cardiac failure^4^, Type 1 diabetes^5^ and retinal degeneration^6^. Beyond therapeutic applications, hiPSCs serve as powerful platforms for disease modeling, allowing the investigation of genetic and multifactorial pathologies in a human cellular context, as well as for substance screening and toxicity testing in drug development, where they provide more predictive and ethically favorable alternatives to animal models.

In the early phase of pluripotent stem cell research, human embryonic stem cells and later hiPSCs were routinely maintained on feeder layers, most commonly mitotically inactivated mouse embryonic fibroblasts (MEFs), in combination with serum-containing or partially undefined media^7^. While these systems supported pluripotency, they suffered from substantial limitations, including xenogeneic contamination, batch-to-batch variability, and poor reproducibility^8^. To address these challenges and enable translational and clinical applications, feeder-free culture systems were developed, incorporating chemically defined media and extracellular matrix (ECM) surrogates that better support standardized, scalable, and regulatory-compliant hiPSC culture.

Among feeder-free culture substrates, Matrigel^TM^ has emerged as the most widely used matrix for hiPSC maintenance and differentiation. Matrigel^TM^ is a solubilized basement membrane extract (BME) derived from Engelbreth–Holm–Swarm (EHS) mouse sarcoma and consists primarily of laminin, collagen IV, nidogen/entactin, and heparan sulfate proteoglycans, along with residual growth factors^9–11^. Its complex and biologically active composition supports robust hiPSC adhesion, survival, and pluripotency, and it is compatible with a wide range of downstream differentiation protocols. However, Matrigel^TM^ presents notable disadvantages, including its undefined and variable composition, lot-to-lot variability, tumor-derived origin, and xenogeneic nature, which limit reproducibility, mechanistic interpretation, and suitability for clinical translation^12^.

To overcome these limitations, defined ECM coatings based on purified or recombinant human proteins have been introduced as alternatives to Matrigel^TM^. Recombinant laminin isoforms such as laminin-511^13,14^ and laminin-521/E-Cadherin^14,15^, as well as vitronectin^16^ and fibronectin-based^17^ substrates, enable precise engagement of integrin-mediated adhesion pathways that support hiPSC self-renewal under fully defined and xeno-free conditions. These systems offer improved reproducibility, reduced biological variability, and enhanced compatibility with good manufacturing practice (GMP). Nevertheless, their use may be associated with higher costs, sensitivity to culture conditions, and, in some cases, reduced versatility across diverse differentiation protocols compared to Matrigel^TM^.

In parallel, fully synthetic matrices have been developed to provide chemically defined and highly tunable environments for hiPSC culture. Early synthetic systems were predominantly based on self-assembled monolayers (SAMs) functionalized with defined peptide sequences^18–21^. Although these substrates have shown promising results with regard to stem cell attachment, maintenance, and differentiation, their suitability for the long-term cultivation of iPSCs remains insufficiently evaluated, as their efficacy has only been demonstrated for selected cell lines. Despite the fact that the Synthemax® substrate based on a Vitronectin-derived synthetic peptide^21^ has already been commercialized, its broad application is not yet achieved. In addition to this, fully synthetic, polymeric substrates, such as MVE-alt-MA^22^, PGC2^23^, and HPhMA-co-HEMA^24^, have been shown to support the culture of both hESCs and hiPSCs. However, sufficient validation for their broader use is still lacking, and their application therefore remains largely confined to academic research settings.

Recently, faCellitate developed a novel fully synthetic, chemically defined ECM-mimicking coating named BIOINSTRUCT^TM^, resembling important features essential for Stem Cell culture. In this study, we investigated whether BIOINSTRUCT^TM^ is suitable for the maintenance of hiPSCs as well as their subsequent differentiation. To this purpose, we cultured IMR 90-4 cells, a standard hiPSC line widely used in basic stem cell research on BIOINSTRUCT^TM^ for 10 passages and compared their stem cell phenotype to hiPSC cultured on Matrigel^TM^. Furthermore, we investigated whether culture on BIOINSTRUCT^TM^ enables multilineage differentiation of hiPSCs and is compatible with Matrigel^TM^-based differentiation protocols by subjecting HIHCNi008-A, ACTN2-GFP, and UKKI-011-A to neuronal, cardiac and pancreatic endocrine differentiation respectively.

## Results

### BIOINSTRUCT^TM^-coating preserves key hiPSC characteristics over long-term culture

To assess the suitability of BIOINSTRUCT^TM^ for the maintenance culture of stem cells, we cultured hiPSCs for 10 passages on BIOINSTRUCT^TM^ (Fig. 1a) and investigated cellular morphology, doubling rate, stem cell marker expression as well as chromosomal integrity. hiPSCs cultured on Matrigel^TM^ served as a reference.

**Figure 1.**
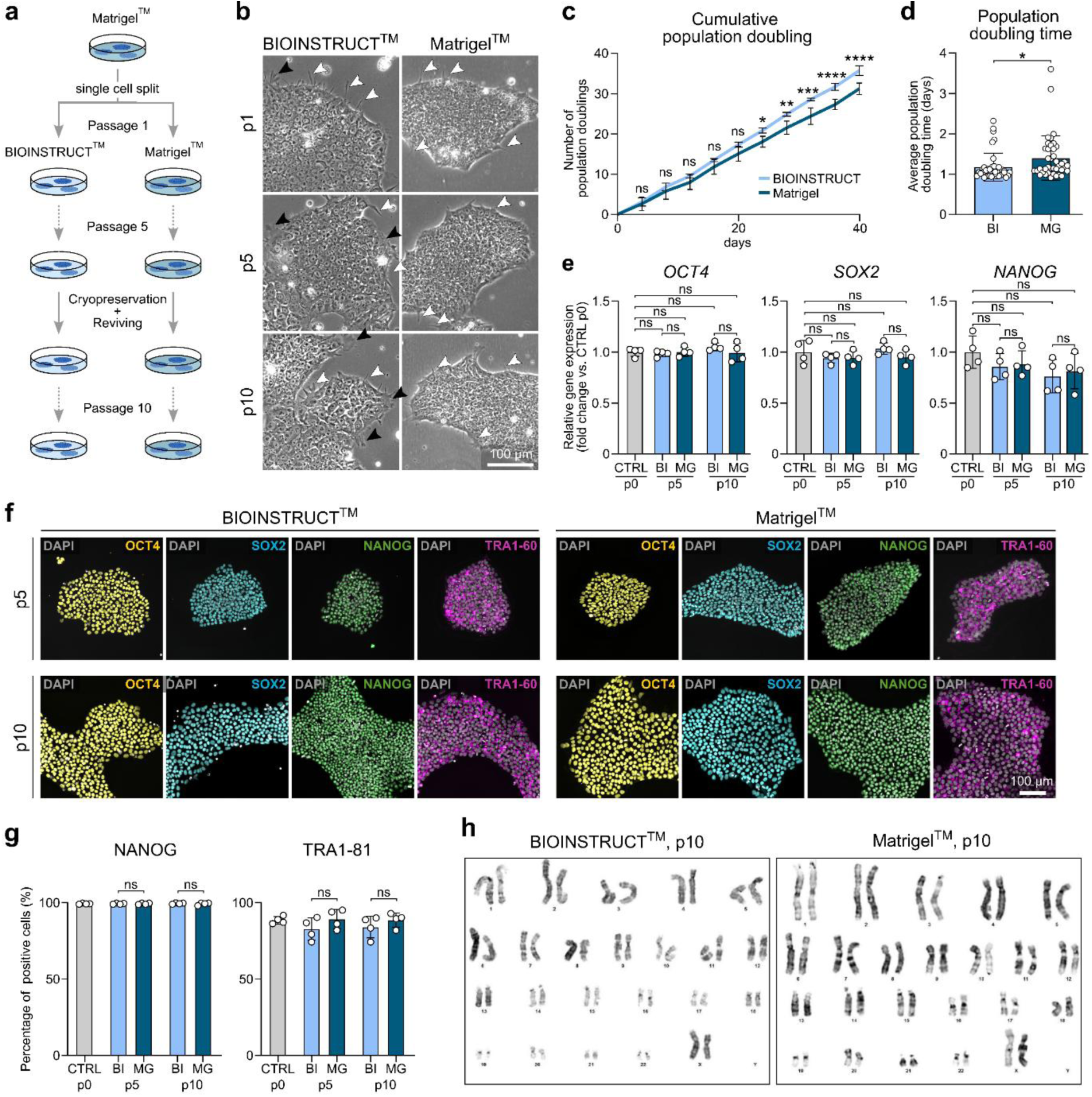
hiPSC maintenance on BIOINSTRUCT^TM^ in comparison to Matrigel^TM^. **a** Graphical illustration of the experimental set up. IMR90-4 hiPSCs were cultured on BIOINSTRUCT^TM^- or Matrigel^TM^-coated plates for 10 passages with single cell passaging every four days and intermediate cryopreservation at passage 5. **b** Brightfield images of hiPSCs cultured on BIOINSTRUCT^TM^ or Matrigel^TM^ at passage 1, 5 and 10 after start of the experiment. **c** Cumulative population doubling of hiPSCs cultured on BIOINSTRUCT^TM^ or Matrigel^TM^ given as the number of population doublings over culture days. Data represent mean ± SD (n=4 independent experiments) for each passage. Statistical analysis was performed using two-way ANOVA with time and surface coating as factors, followed by Sidak’s multiple comparisons test. ns (not significant) P > 0.05, *P < 0.05, **P < 0.01, ***P < 0.005, P**** < 0.001. **d** Average population doubling time of hiPSCs cultured on BIOINSTRUCT^TM^ or Matrigel^TM^. Data represent mean ± SD. Data points represent the population doubling time of single passages (n=40 from 4 independent experiments). Statistical analysis was performed using an unpaired two-tailed t-test. *P < 0.05. **e** Gene expression of the core pluripotency genes *OCT4*, *SOX2*, and *NANOG* of hiPSCs cultured on BIOINSTRUCT^TM^ or Matrigel^TM^ at passage 5 and 10 normalized to the expression at the start of the experiments (passage 0). Data represent mean ± SD (n=4 independent experiments). Statistical analysis was performed using one-way ANOVA followed by Sidak’s multiple comparisons test. ns (not significant) P > 0.05. **f** Representative immunofluorescent images of hiPSCs cultured on BIOINSTRUCT^TM^ or Matrigel^TM^ at passage 5 and 10. hiPSCs are stained for OCT4 (yellow), SOX2 (cyan), NANOG (green) and TRA1-60 (magenta). hiPSCs are counterstained with DAPI (grey). **g** Bar graphs of flow cytometry analysis, showing the number of NANOG-positive and TRA1-81-positive hiPSCs at passage 5 and 10 of culture on BIOINSTRUCT^TM^ or Matrigel^TM^, as well as before the start of the experiment (passage 0). Data represent mean ± SD (n=4 independent experiments). Statistical analysis was performed using one-way ANOVA followed by Sidak’s multiple comparisons test. ns (not significant) P > 0.05. **h** Karyogramm of hiPSC at passage 10 of culture BIOINSTRUCT^TM^ or Matrigel^TM^. For the karyogramm hiPSCs from two experiments were pooled and analyzed. Abbreviations: BI: BIOINSTRUCT^TM^, MG: Matrigel^TM^, p: passage. An overview of hiPSC morphology during all 10 passages is displayed in Supplementary Figure 1.

hiPSCs cultured on BIOINSTRUCT^TM^ retained a characteristic hiPSC-like morphology over 10 passages, displaying round, densely packed colonies with a high nucleus-to-cytoplasm ratio (Fig. 1b). Compared to Matrigel^TM^, hiPSCs maintained on BIOINSTRUCT^TM^ appeared slightly larger with a marginally reduced nucleus-to-cytoplasm ratio, an effect that was particularly evident at the colony edges. Furthermore, cells cultured on BIOINSTRUCT^TM^-coated plates exhibited an increased number of pseudopodia at the colony borders (white and black arrowheads). In addition to thin cellular extensions (white arrowheads), which were observed at lower frequency on Matrigel^TM^, hiPSCs grown on BIOINSTRUCT^TM^ formed broad, flattened pseudopodia (black arrowheads) that were rarely detected under Matrigel^TM^ conditions. A detailed characterization of cell morphology across all passages is provided in Supplemental Figure 1.

hiPSCs cultured on BIOINSTRUCT^TM^ exhibited a slightly higher expansion rate compared to those cultured on Matrigel^TM^. Over 10 passages, hiPSCs maintained on BIOINSTRUCT^TM^ achieved 35.8 ± 1.2 population doublings within 40 days, compared to 31.3 ± 1.4 population doublings on Matrigel^TM^ (Fig. 1c). This corresponded to average population doubling times of 1.18 ± 0.35 days and 1.41 ± 0.55 days, respectively (Fig. 1d).

Following confirmation of the maintenance of hiPSC-like morphology and expansion capacity, we next assessed the expression of pluripotency-associated markers. hiPSCs cultured on BIOINSTRUCT^TM^ retained expression of *OCT4*, *SOX2*, and *NANOG* over 10 passages, comparable to cells cultured on Matrigel^TM^ (Fig. 1e). While *OCT4* and *SOX2* expression remained stable throughout the culture period, *NANOG* expression showed a slight decrease over time under both conditions (BIOINSTRUCT^TM^ p5: 0.85 ± 0.13-fold, p10: 0.76 ± 0.17-fold; Matrigel^TM^ p5: 0.89 ± 0.13-fold; p10: 0.81 ± 0.18-fold), although this was not statistically significant. Immunofluorescence staining further confirmed the expression of OCT4, SOX2, NANOG, and TRA1-60 in hiPSCs cultured on BIOINSTRUCT^TM^ (Fig. 1f). The proportion of NANOG⁺ and TRA-1-81⁺ cells remained unchanged during long-term culture on BIOINSTRUCT^TM^ (99.5 ± 0.4% and 84.2 ± 7.2%, respectively) and was comparable to Matrigel^TM^ conditions (99.3 ± 0.8% and 88.7 ± 4.7%) (Fig. 1g). Karyotype analysis demonstrated a normal chromosomal profile, confirming the genomic stability of hiPSCs maintained on BIOINSTRUCT^TM^ (Fig. 1h).

Tests for the suitability of BIOINSTRUCT^TM^ for the maintenance culture were replicated with a second cell line (Supplemental Figure 2a-f). The collected data confirm the retention of morphology, stem cell marker expression, and genomic integrity of BIONi010B cells during the experimental period of 10 passages, providing a first evidence for a cell-line independent suitability of BIOINSTRUCT^TM^ for stem cell maintenance culture.

Taken together, these data demonstrate that culture on the synthetic ECM surrogate BIOINSTRUCT^TM^ preserves key hiPSC characteristics and supports stable hiPSC expansion comparable to Matrigel^TM^.

### BIOINSTRUCT^TM^-coating preserves hiPSC pluripotency and enables multilineage differentiation

After confirming the maintenance of a characteristic hiPSC phenotype on BIOINSTRUCT^TM^, we next investigated whether hiPSCs cultured on BIOINSTRUCT^TM^ for 10 passages retained their potential to generate derivatives of all three germ layers. Furthermore, we assessed whether BIOINSTRUCT^TM^ could support multilineage differentiation when used as a coating substrate during the differentiation process. To this end, hiPSCs cultured on BIOINSTRUCT^TM^ or Matrigel^TM^ for 11 passages were first aggregated into embryoid bodies (EBs) and subsequently subjected to spontaneous differentiation by plating EBs onto BIOINSTRUCT^TM^- or Matrigel^TM^ -coated plates (Fig. 2a). EBs generated from hiPSCs maintained on either BIOINSTRUCT^TM^ or Matrigel^TM^ successfully attached and spread on both coating substrates. Continuous spontaneous differentiation resulted in the emergence of cell populations displaying distinct morphologies at days 16 and 26 (Fig. 2b), indicating the generation of diverse cell types derived from different developmental lineages.

**Figure 2.**
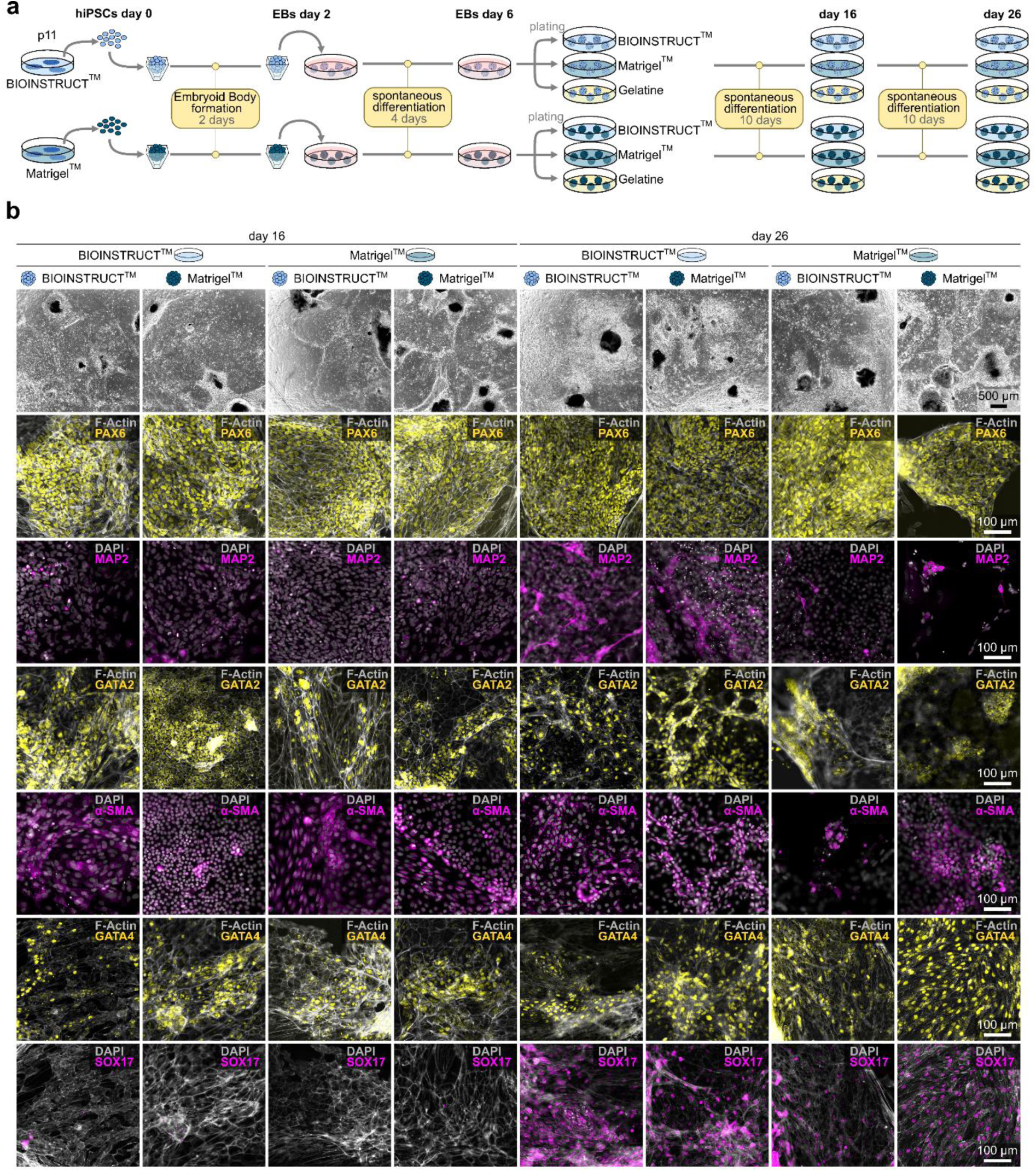
Spontaneous differentiation on BIOINSTRUCT^TM^ and Matrigel^TM^. **a** Scheme of the experimental set-up. hiPSCs cultured on BIOINSTRUCT^TM^ or Matrigel^TM^ for 11 passages (hiPSCs d0), were used to form Embroyid bodies (EBs day 2). Spontaneous differentiation was induced in suspension for 4 days (EBs day 6), before EBs were plated on gelatine-, BIOINSTRUCT^TM^- or Matrigel^TM^-coated plates and cultured in spontaneous differentiation medium for another 10 (day 16) and 20 days (day 26). **b** Representative images of EBs originated from p11-BIOINSTRUCT^TM^ or p11-Matrigel^TM^-cultured hiPSCs, plated on BIOINSTRUCT^TM^ or Matrigel^TM^ and analyzed on day 16 or day 26 of culture. Shown are representative brightfield images, demonstrating the formation of cell populations with distinct cellular morphologies (panel 1), as well as immunofluorescent images of staining against PAX6 (yellow), MAP2 (magenta), GATA2 (yellow), α-SMA (magenta), GATA4 (yellow), and SOX17 (magenta). Depending on the nuclear or cytoplasmic localization of the respective antigen, counterstaining with Phalloidin to visualize the F-Actin cytoskeleton (grey) or DAPI to stain the cell nuclei is shown.

To further confirm lineage specification, spontaneously differentiated cultures were analyzed for the expression of markers associated with the three germ layers. The presence of PAX6, an essential regulator of neuroectodermal development, together with MAP2, a marker of neuronal maturation, indicated the generation of ectoderm-derived structures under all experimental conditions. While PAX6 was already detectable at day 16 of spontaneous differentiation, MAP2-positive cells exhibiting neuronal morphology were observed only at later stages of differentiation. Cells differentiated under all conditions further exhibited GATA2, a marker associated with hematopoietic, endothelial, neural crest, and extraembryonic lineages, as well as αSMA, a marker of mesenchymal maturation, demonstrating the induction of mesodermal lineage differentiation. Endodermal commitment was confirmed by the presence of GATA4, which is involved in endodermal and mesodermal developmental programs, and SOX17, a marker of definitive endoderm formation. While GATA4 was detected at both days 16 and 26 of spontaneous differentiation, SOX17 was observed only at the later time point.

Preservation of pluripotency could also be confirmed for BIONi010B cells after 10 passages culture on BIOINSTRUCT^TM^. Trilineage differentiation resulted in the formation of derivatives of all three germ layers as confirmed by the presence of Nestin (ectoderm), Brachyury (mesoderm), and SOX17 (endoderm) (Supplemental Figure 2g).

Taken together, these findings indicate that hiPSCs maintained on BIOINSTRUCT^TM^ retain their intrinsic capacity to initiate differentiation programs associated with all three germ layers. The data further demonstrate that BIOINSTRUCT^TM^ enables multilineage differentiation when used as a culture substrate during spontaneous differentiation.

### BIOINSTRUCT^TM^-coating does not affect developmental capacity in comparison to Matrigel^TM^

After demonstrating that BIOINSTRUCT^TM^ supports the formation of derivatives from all three germ layers, we next investigated whether long-term maintenance on BIOINSTRUCT^TM^ or the use of BIOINSTRUCT^TM^ as a differentiation substrate introduces lineage bias during spontaneous differentiation. To quantitatively assess the multilineage differentiation capacity of hiPSCs, we determined germ layer scores based on the standardized approach introduced by D’Antonio et al.^25^ using an adapted target gene panel. To distinguish potential effects arising from the maintenance phase from those caused by the differentiation substrate itself, EBs generated from hiPSCs maintained on BIOINSTRUCT^TM^ or Matrigel^TM^ were differentiated on BIOINSTRUCT^TM^, Matrigel^TM^, or gelatin-coated surfaces (Fig. 2a).

Germ layer score analysis revealed comparable ectodermal and mesodermal differentiation scores between BIOINSTRUCT^TM^- and Matrigel^TM^-maintained hiPSCs under all tested differentiation conditions (Fig. 3a). Endodermal scores were moderately reduced in cells derived from BIOINSTRUCT^TM^ maintenance cultures at day 16 and remained slightly decreased at day 26, suggesting a potential delay or altered kinetics of endoderm-associated differentiation. Furthermore, we detected an increased ectodermal score for cells from Matrigel^TM^ maintenance culture plated on BIOINSTRUCT^TM^.

**Figure 3.**
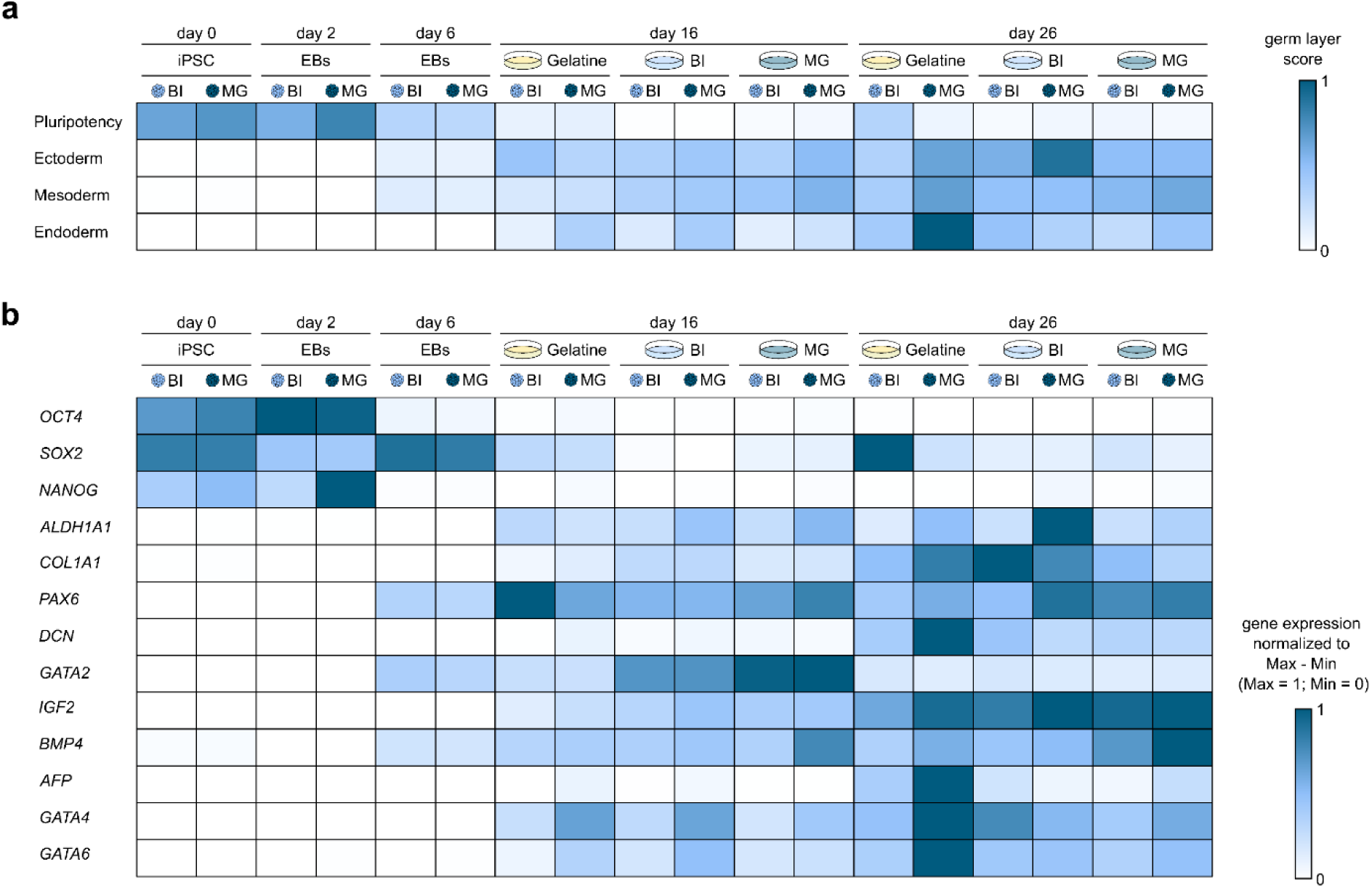
Germ-layer associated gene expression of spontaneous differentiated hiPSCs on BIOINSTRUCT^TM^, Matrigel^TM^ and Gelatine. **a** Heat map showing the Germ Layer Scores for each sample. The Germ Layer Score was calculated as the mean score of genes associated to the same germ layer according to D’antonio et al^24^ (n ≥ 3 independent experiments). Shown is the expression for hiPSCs (d0), after formation of EBs (d2), following induction of spontaneous differentiation in suspension (EBs d6), and after plating of EBs and continued differentiation on gelatin-, BIOINSTRUCT^TM^- or Matrigel^TM^-coated plates at day 16 and day 26. **b** Expression of the individual genes used to determine the germ layer score shown as a heat map. Gene expression of each sample has been normalized to the range of minimal (=0) and maximal (=1) expression across all samples of one gene (n ≥ 3 independent experiments). Bar graphs of the gene expression data including statistical analyses are displayed in Supplementary figures 3 and 4.

To further evaluate lineage-associated transcriptional changes independently of predefined germ layer assignments, individual marker expression patterns were analyzed (Fig. 3b). Overall, expression profiles were comparable between BIOINSTRUCT^TM^ and Matrigel^TM^ conditions, with only minor differences observed. When differentiation was performed on gelatin, cells derived from BIOINSTRUCT^TM^-maintained hiPSCs exhibited generally lower expression levels across several lineage-associated markers at day 26 compared to cells derived from Matrigel^TM^ maintenance cultures. In particular, reduced expression of the endoderm-associated markers *GATA4* and *GATAc* at day 16, as well as *AFP* at day 26 was observed, which explains the reduction in endodermal scores in BIOINSTRUCT^TM^-maintained cultures. Comparison of the differentiation substrates indicated a substrate-dependent trend for *GATA2* expression at day 16, with lowest expression in gelatin-coated cultures, intermediate expression in BIOINSTRUCT™-coated cultures, and highest expression in Matrigel^TM^-coated cultures. However, this difference was not reflected by an overall alteration of mesodermal differentiation scores. *COL1A1* expression was slightly increased in cells plated on BIOINSTRUCT^TM^ for spontaneous differentiation, whereas *PAXc* was slightly elevated in cultures plated on Matrigel^TM^. *ALDH1A1* exhibited increased expression in cells originating from Matrigel^TM^-maintenance cultures at both, day 16 and day 26. Although all three ectoderm associated genes exhibited a substrate-dependent differential expression, this did not result in a shift of the ectodermal score. We further detected an increased *SOX2* expression in BIOINSTRUCT^TM^ -maintained cultures plated on Gelatine at day 26, which was due to an outlier of a single experiment (Supplemental Fig. 3) and resulted in an increase of the pluripotency score in this condition. Individual expression values of the analyzed genes as well as statistical analysis are displayed in Supplemental Figure 3 and 4.

**Figure 4.**
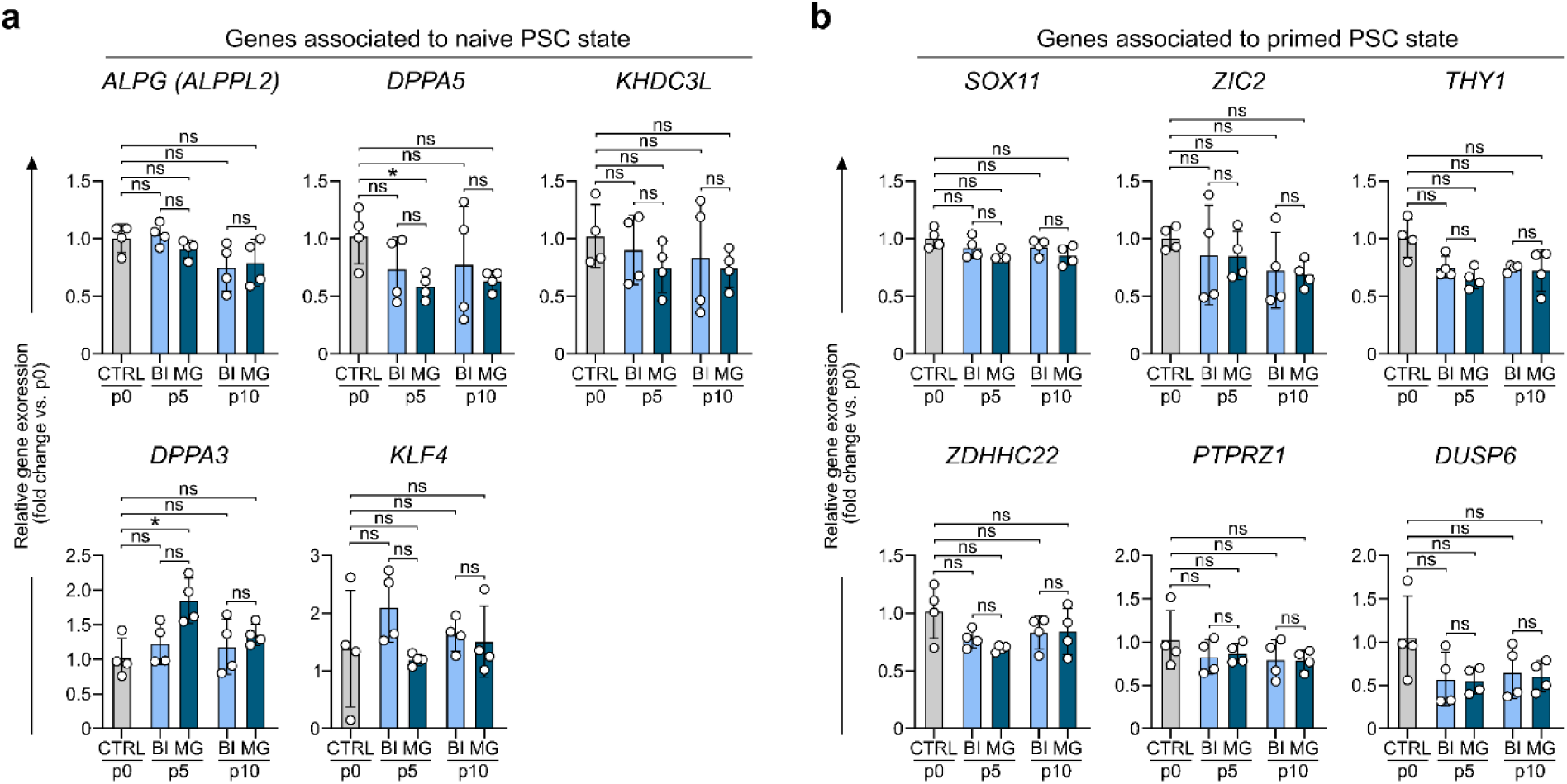
Determination of naïve and primed ground state of hiPSCs cultured on BIOINSTRUCT^TM^. a, **b** Gene expression of genes associated with a naïve (*ALPG* (*ALPPL2*), *DPPA5*, *KHDC3L*, *DPPA3*, *KLF4*) and primed (*SOX11*, *ZIC2*, *THY1*, *ZDHHC22*, *PTPRZ1*, *DUSPc*) pluripotency state. Gene expression is shown for hiPSCs at the beginning of the experiment (CTRL p0) as well as at p5 and p10 of culture on BIOINSTRUCT^TM^ or Matrigel^TM^. Gene expression is displayed as fold change relative to CTRL p0. Data represent mean ± SD (n=4 independent experiments). Statistical analysis was performed using one-way ANOVA followed by Sidak’s multiple comparisons test. ns (not significant) P > 0.05, *P < 0.05.

Given the differences in early germ layer gene expression in cells maintained on BIOINSTRUCT^TM^ compared to hiPSCs maintained on Matrigel^TM^, particularly regarding the endoderm lineage, we were interested to learn whether these arose from a potential shift in the pluripotency state. We therefore investigated the ground state of hiPSCs cultured on BIOINSTRUCT^TM^ and Matrigel^TM^ by analyzing the expression of genes associated with a naïve and primed stem cell state^26,27^. hiPSCs cultured on BIOINSTRUCT^TM^ and Matrigel^TM^ exhibited similar expression profiles of genes associated with a naïve (Fig. 4a) and primed state (Fig. 4b). Significant changes in expression were only observed for the naïve PSC state markers *DPPA5* (0.59-fold ± 0.09) and *DPPA3* (1.85-fold ± 0.28) in hiPSCs cultured on Matrigel^TM^ for 5 passages compared to the hiPSCs at the beginning of the experiment (Fig. 4a). In summary, the data indicate that maintenance culture on BIOINSTRUCT^TM^ did not affect the ground state of hiPSC in comparison to Matrigel^TM^.

Taken together, the overall comparable germ layer scores and lineage-associated expression profiles between BIOINSTRUCT^TM^ and Matrigel^TM^ conditions indicate that BIOINSTRUCT^TM^ does not induce a lineage bias during long-term hiPSC maintenance or spontaneous differentiation. Furthermore, the data show that BIOINSTRUCT^TM^ supports multilineage differentiation when used as a culture substrate during differentiation.

### BIOINSTRUCT^TM^-coating supports the formation of hiPSC-derived neuronal cells, cardiomyocytes and pancreatic endocrine cell types

After showing that BIOINSTRUCT^TM^-coating enables maintenance and undirected differentiation of hiPSCs comparable to Matrigel^TM^, we next investigated whether BIOINSTRUCT^TM^ is compatible with established Matrigel^TM^-based differentiation protocols for the generation of diverse cell types. To this end, hiPSCs of three different cell lines were subjected to directed differentiation towards neuronal, cardiac, and pancreatic endocrine lineages, covering ectodermal, mesodermal, and endodermal lineage differentiation routes, respectively.

To investigate the compatibility with neuronal differentiation, hiPSCs were differentiated to cortical neurons on BIOINSTRUCT^TM^ - or Matrigel^TM^-coated plates according to previously reported protocols based on dual-SMAD inhibition (Fig. 5a)^28–30^. At the final replating step, surfaces were pre-coated with poly-L-ornithine (1h at RT) prior to BIOINSTRUCT^TM^ and Matrigel^TM^ coating, as established in the Matrigel^TM^ based workflow. BIOINSTRUCT^TM^ without poly-ornithine pre-coating did not support terminal differentiation reliably, as cells detached before day 36 of the protocol. On both coatings a similar morphological progression of differentiating cultures could be observed as show for day 19 and 36 (Fig. 5b). At day 36, immunocytochemical staining revealed robust TUJ1 and CTIP2 signals in cultures differentiated on BIOINSTRUCT^TM^ and Matrigel^TM^, whereas GFAP signal was not detected in either condition (Fig. 5c). Neuronal morphology was similar on both coatings. RT-qPCR analysis of neuronal and cortical markers (*TUBB3*, *MAP2*, *BCL11B*, *TBR1*, *SLC17Ac*, *MAPT*, *FOXG1*, *SOX2*, and *SYN1*) did not reveal statistically significant differences in expression between coating conditions (Fig. 5d). In line with the immunofluorescent staining, no expression of the astrocytic marker *GFAP* was detected in any sample (data not shown), indicating pure neuronal differentiations. To test whether cortical neurons generated on BIOINSTRUCT^TM^ form functional networks, neurons were replated on day 26 onto multiwell microelectrode arrays coated with poly-L-ornithine and BIOINSTRUCT^TM^ or Matrigel^TM^, co-cultured with primary mouse astrocytes, and recorded from day 43 to day 87 (Fig. 5e). On both coatings, spontaneous spiking was detected from the first recording onwards and increased over time. Network bursts emerged on both coatings, on BIOINSTRUCT^TM^ with a delayed onset from day 66. By the late phase spike frequency had reached similar levels. Representative spike trains at day 87 showed recurring network bursts synchronized across electrodes on both coatings (Fig. 5f). Taken together, these findings indicate that BIOINSTRUCT^TM^ supports the formation of spontaneously active, synchronized neuronal networks using an established Matrigel^TM^ -based differentiation protocol.

**Figure 5.**
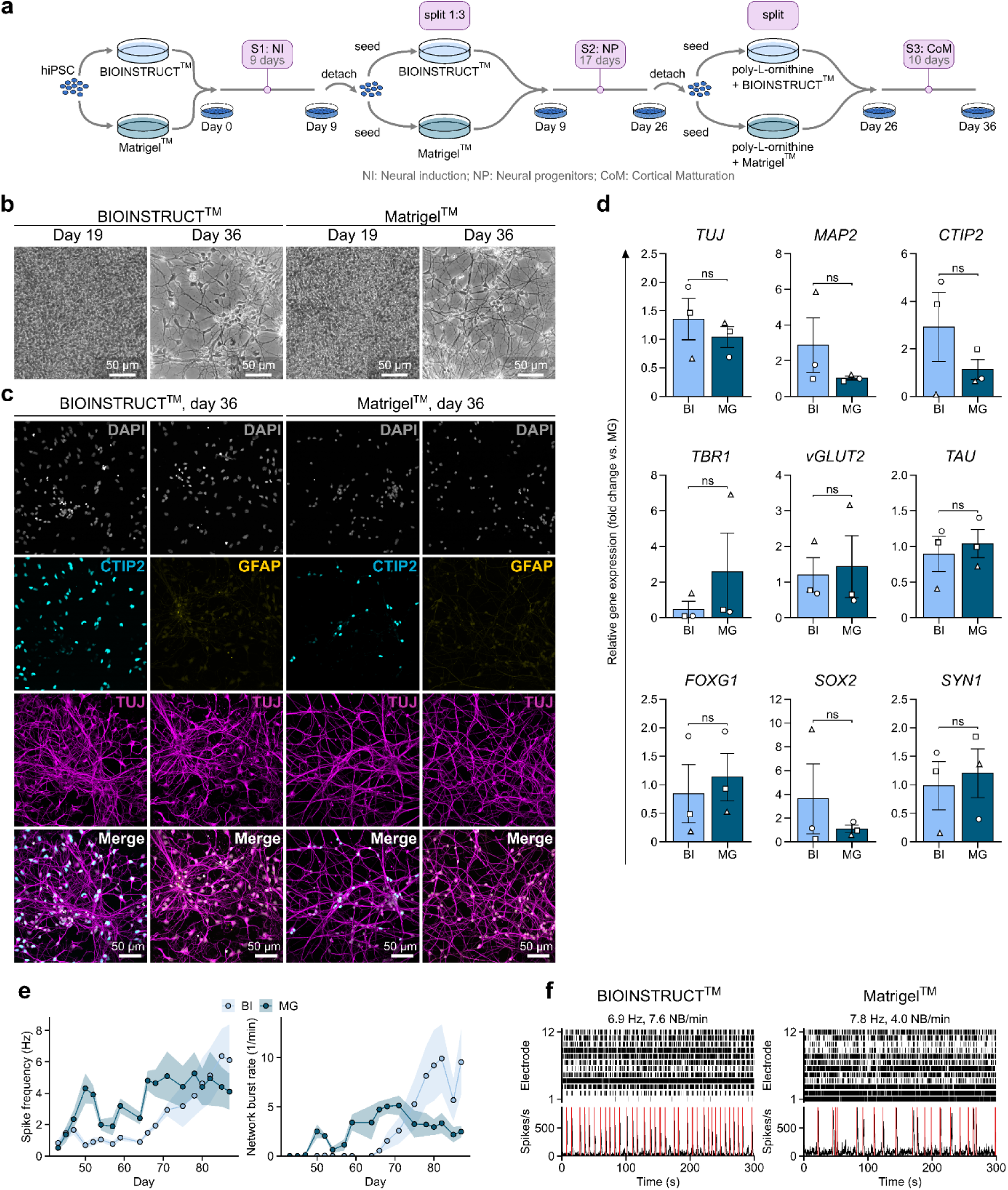
Cortical neurons differentiated on BIOINSTRUCT^TM^ versus Matrigel^TM^. **a** Graphical illustration of the experimental workflow. hiPSCs were cultivated on BIOINSTRUCT^TM^ or Matrigel^TM^ -coated plates and seeded on the respective matrix in high density (d0) followed by neural induction (NI), neural precursor proliferation (NP) and cortical maturation (CoM). **b** Representative brightfield images of neural precursors (day 19) and mature cortical neurons (day 36) on BIOINSTRUCT^TM^ and Matrigel^TM^, respectively. **c** Exemplary confocal images of immunostainings for the pan-neuronal marker β-III-tubulin (TUJ, magenta)), the cortical marker CTIP2 (cyan) and the astrocytic marker GFAP (yellow). Shown are maximum intensity projections of confocal z-stacks, nuclei counterstained with DAPI (grey). **d** RT-qPCR analysis of neuronal marker expression in three independent differentiations (individual symbols). Expression values were normalized to the respective Matrigel^TM^ condition. Data represent mean ± SEM (n=3 independent experiments). Statistical analysis was performed using two-sided Mann-Whitney U test. ns (not significant) P > 0.05. **e** Spontaneous network activity of cortical neurons co-cultured with primary mouse astrocytes on multiwell microelectrode arrays (MEA) coated with poly-L-ornithine and BIOINSTRUCT or Matrigel, recorded from day 43 to day 87. Shown are spike frequency per active electrode and network burst rate. Data represent mean ± SEM (n=6 wells from one differentiation, 18 recordings). Wells without network bursts were included as 0. **f** Representative spike trains of one BIOINSTRUCT^TM^ and one Matrigel^TM^ well at day 87, shown over the full 5 min recording as raster plots of the 12 electrodes per well with the summed spike rate in 100 ms bins below (red lines mark network burst onsets). Values above the rasters give the spike frequency and network burst rate of the respective well.

Next, we investigated the suitability of BIOINSTRUCT^TM^ for the generation of cardiac cells. For this purpose, ventricular cardiomyocytes were generated using the method laid out by Dark et al.^31^. Cells were replated once during the differentiation, and cultured to day 30 (Fig. 6a), at which point they were analyzed for gene expression, ACTN2 (alpha-actinin-2) expression, morphology and contraction. Cells differentiated on either surface produced high-purity hiPSC-derived cardiomyocytes (hiPSC-CMs), as percentage of cells differentiated on either BIOINSTRUCT^TM^ or Matrigel^TM^ was 99.79 ± 0.1% and 99.82 ± 0.17, respectively (Fig. 6b,c). There was no significant difference in average ACTN2 expression per cell (Fig. 6d). Nucleus size was 2% larger for cells differentiated on Matrigel^TM^, with an average diameter of 14.88 ± 0.17 µm versus 14.5 ± 0.14 µm (Fig. 6e). There was no difference in nucleus circularity, which was 0.6 ± 0.01 arbitrary units for both conditions (Fig. 6f). The expression of key cardiac structural genes (*TTN*, *MYHc*, *MYH7*) and cardiac calcium channels (*RYR2*, *ATP2A2*) was measured using RT-qPCR, relative to BIOINSTRUCT^TM^ expression and normalized using *GAPDH* (Fig. 6g). There was no difference in *TTN* expression, however *MYHc* and *MYH7* were 1.76 ± 0.26 and 1.61 ± 0.32-fold higher in cells differentiated on Matrigel^TM^. *RYR2* and *ATP2A2* were expressed lower, at 0.67 ± 0.17 and 0.57 ± 0.12-fold, respectively. Spontaneous contraction was measured by recording the cells under normal culture conditions (37°C, 5% CO_2_, atmospheric O_2_) followed by analysis using MUSCLEMOTION^32^. This software uses pixel change to detect movement and generate a graphical trace representing each contraction event (or peak). Contraction duration refers to the number of milliseconds between the start and end of the peak. Contraction amplitude refers to the distance between the base and tip of the peak, a pseudo-measurement of contractile force, where greater pixel movement results in a greater contraction amplitude. Contraction frequency is calculated from the number of peaks divided by the total time per recording and converted into Hz (peaks per second). Contraction frequency was slower for hiPSC-CMs on Matrigel^TM^-coated plates, at 0.25 ± 0.06 Hz compared to 0.49 ± 0.07 Hz (Fig. 6i). Contraction amplitude was 2.3x greater for cells generated on Matrigel^TM^ (Fig. 6j), while contraction duration of Matrigel^TM^-generated cells was 1353 ± 258 ms, which was 404 ms longer than cells on BIOINSTRUCT^TM^ (Fig. 6k). Altogether, the data demonstrate that hiPSCs cultured on BIOINSTRUCT^TM^ were capable of generating functional cardiomyocytes, although data suggest there may be some differences in contraction and morphology, possibly due to differences in attachment level between the cells and the matrix, or differences in signaling from the matrix during development.

**Figure 6.**
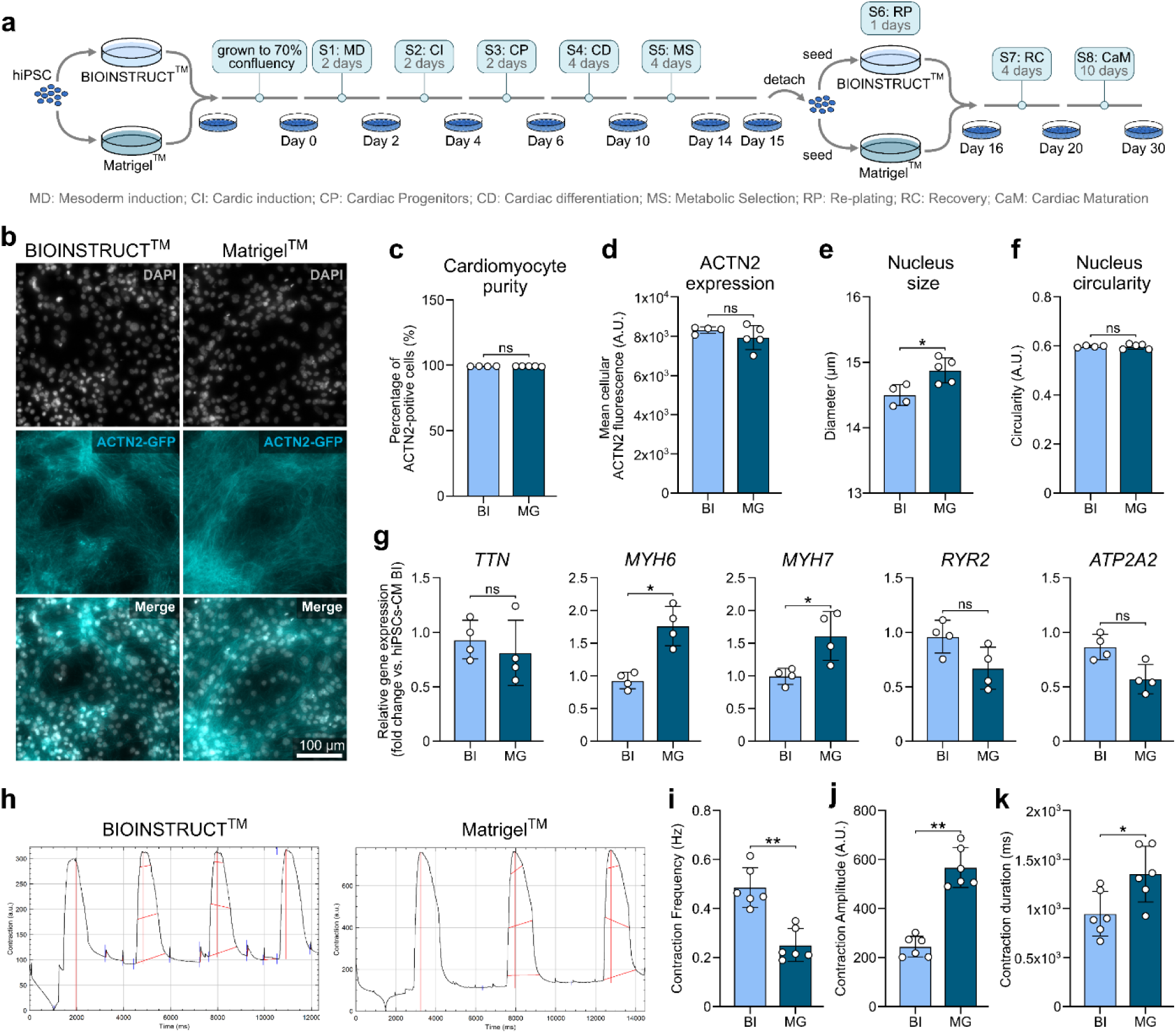
Functional cardiomyocytes differentiated on BIOINSTRUCT^TM^. **a** Graphical illustration of the experimental procedure. ACTN2-GFP hiPSC were seeded on BIOINSTRUCT^TM^- or Matrigel^TM^-coated plates and differentiated towards cardiac myocytes via the mesodermal lineage following a protocol adapted from Dark et al.^30^. **b** Representative fluorescent images of ACTN2-GFP hiPSC-CMs (cyan) at day 30 of differentiation on BIOINSTRUCT^TM^- or Matrigel^TM^-coated plates, fixed and stained with DAPI (grey). **c-f** Quantitative analysis of fluorescent images showing hiPSC-CMs purity (c), ACTN2 expression (d), nucleus size (e) and (iv) nucleus circularity (f). Data represent mean ± SD (n=4 independent experiments for BI, n=5 for MG). Statistical analysis was performed using Mann-Whitney test. ns (not significant) P > 0.05, *P < 0.05. **g** Gene expression analysis of *TTN*, *MYHc*, *MYH7*, *RYR2* and *ATP2A2* shown as foldchange expression compared to hiPSC-CMs differentiated on BIOINSTRUCT^TM^. Data represent mean ± SD (n=4 independent experiments). Statistical analysis was performed using Mann-Whitney test. ns (not significant) P > 0.05, *P < 0.05. **h** 2D contraction analysis of day 30 hiPSC-CMs differentiated on (i) BIOINSTRUCT^TM^- or (ii) Matrigel^TM^-coated plates. Cells were recorded for 10-20 seconds per recording and contraction traces were generated using MUSCLEMOTION^31^. **i-k** Analysis of the contraction traces to calculate contraction frequency (i), contraction amplitude (j) and contraction duration (k). Data represent mean ± SD (n=6 independent experiments, with 5 recordings per experiment). Statistical analysis was performed using Mann-Whitney test. ns (not significant) P > 0.05, *P < 0.05, ** P < 0.01.

In addition, we tested whether BIOINSTRUCT^TM^ is also compatible with protocols for the generation of hiPSC-derived pancreatic endocrine cells. Directed pancreatic differentiation was performed according to a protocol established by Balboa and Barsby et al^33,34^. Briefly, hiPSCs were differentiated toward pancreatic progenitors during 10 days of adherent culture, followed by detachment at day 11 and continued differentiation as spheroids in suspension culture for an additional 37 days to generate stem-cell-derived islet-like clusters (SC-Islets) (Fig. 7a). Similar to Matrigel^TM^ conditions, hiPSCs seeded onto BIOINSTRUCT^TM^ reached full confluency within 24 h (Fig. 7b). Sequential induction of pancreatic developmental stages on BIOINSTRUCT^TM^ resulted in characteristic morphological changes and spheroid formation comparable to Matrigel^TM^-cultured cells. At the posterior gut tube stage, cultures maintained on BIOINSTRUCT^TM^ displayed a less compact upper cell layer compared to Matrigel^TM^ conditions, suggesting potential differences in the efficiency of subsequent pancreatic endocrine differentiation.

**Figure 7.**
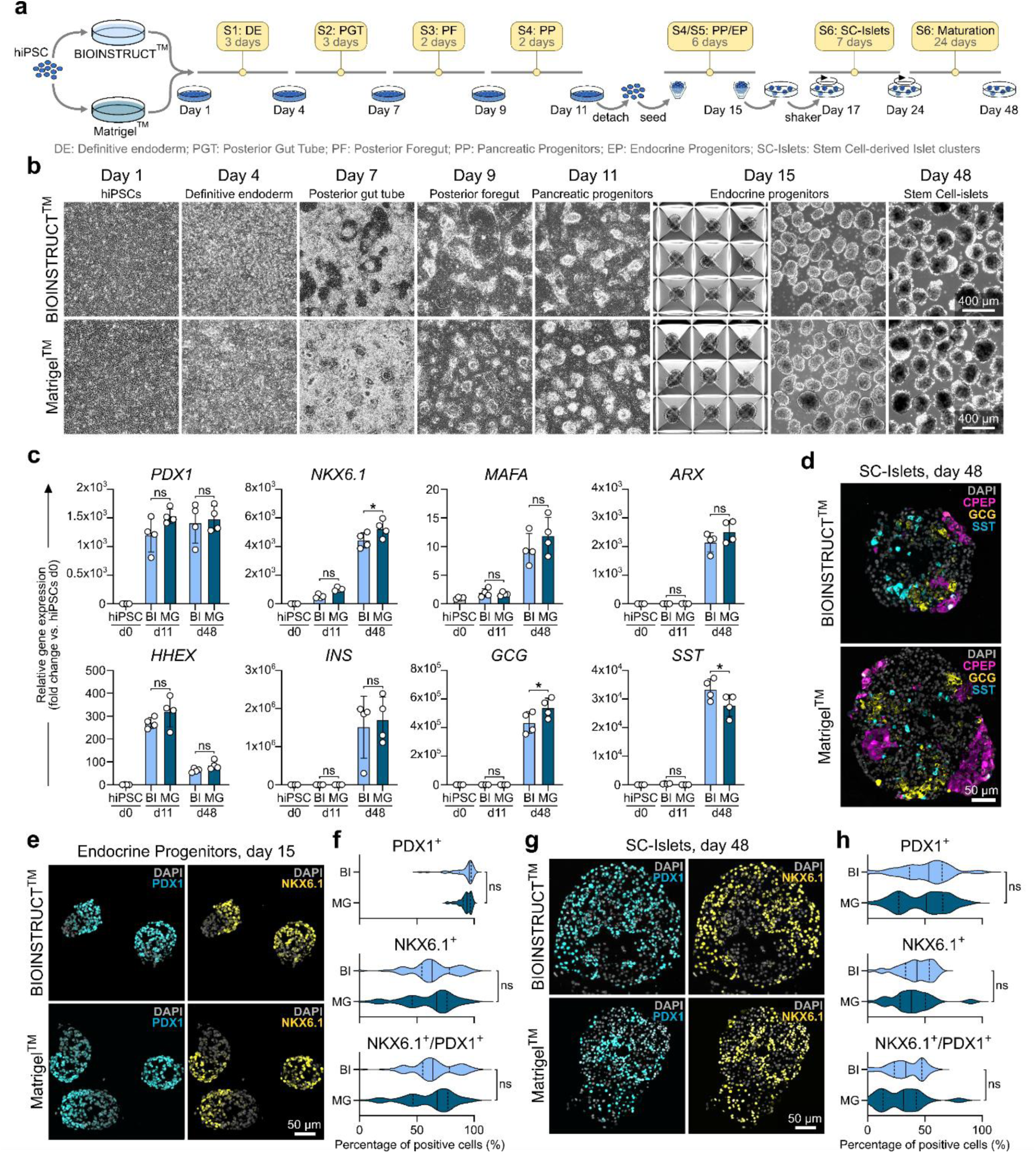
Formation of pancreatic Islet-like cell clusters on BIOINSTRUCT^TM^. **a** Scheme of the experimental set up. UKKI-011A hiPSCs were seeded on BIOINSTRUCT^TM^- or Matrigel^TM^-coated plates and differentiated towards pancreatic islet-like clusters via the endodermal lineage following a protocol developed by Barsby et al. and Balboa et al.^32,33^. **b** Representative brightfield images of different stages of pancreatic differentiation on BIOINSTRUCT^TM^ or Matrigel^TM^. **c** Gene expression analysis of transcription factors hallmarking pancreatic endocrine lineage commitment (*HHEX*, *PDX1*, *NKXc.1*), as well as transcription factors and hormones associated with an α-cell (*ARX*, *GCG*), β-cell (*NKXc.1*, *MAFA*, *INS*), and δ-cell (*HHEX*, *SST*) fate. Gene expression was analyzed at d11 and d48 of differentiation on BIOINSTRUCT^TM^ or Matrigel^TM^ and normalized to expression of hiPSCs (day 0). Data represent mean ± SD (n=4 independent experiments). Statistical analysis was performed using one-way ANOVA followed by Sidak’s multiple comparisons test. ns (not significant) P > 0.05, *P < 0.05. **d** Immunofluorescently labelled crosssection of SC-Islets at day 48 of pancreatic differentiation, generated from cells differentiated on BIOINSTRUCT^TM^ or Matrigel^TM^. Samples are stained for the α-cell hormone GCG (yellow), the β-cell marker CPEP (magenta) and the δ-cell hormone SST (cyan). Samples are counterstained with DAPI. **e-h** immunofluorescent staining of crosssection of spheroids at day 15 and 48 of pancreatic differentiation, induced on either BIOINSTRUCT^TM^ or Matrigel^TM^. Crosssections are immunofluorescently labelled with antibodies against the pancreatic lineage markers PDX1 (cyan) and NKX6.1 (yellow) and counterstained with DAPI (e, g). Violine plots show the ratio of PDX1-positive, NKX6.1-positive, and PDX/NKX6.1 double-positive cells, quantified by automated cell counting (f, h). Violin plots display the median (continuous line) and quartiles (dashed lines). Data are based on the analysis of single spheroids with n=43 for BIOINSTRUCT^TM^ d15, n=72 for Matrigel^TM^ d15, n=73 for BIOINSTRUCT^TM^ d48, and n=36 for Matrigel^TM^ d48 from ≥ 3 independent experiments). Statistical analysis was performed using an unpaired two-tailed t-test. ns (not significant) P > 0.05.

Gene expression analysis revealed that *PDX1*, a master regulator of pancreatic lineage specification, was maintained at comparable levels in cells differentiated on BIOINSTRUCT^TM^ (Fig. 7c). In contrast, *NKXc.1*, a key regulator of endocrine lineage commitment and β-cell maturation, showed a slight reduction at day 48 in BIOINSTRUCT^TM^-derived cultures (d48: 4.4 x 10^3^ ± 0.5 x 10^3^-fold) compared to Matrigel^TM^ (d48: 5.3 x 10^3^ ± 0.6 x 10^3^-fold). Despite this difference, expression of the β-cell-associated markers *MAFA* and *INS* remained unchanged between both conditions. Interestingly, SC-Islets generated on BIOINSTRUCT^TM^ exhibited slightly reduced expression of the α-cell hormone *GCG* at day 48 (4.4 x10^5^ ± 0.8 x10^5^-fold) compared to Matrigel^TM^ (5.4 x10^5^ ± 0.7 x10^5^-fold), accompanied by a modest increase in expression of the δ-cell hormone *SST* (BIOINSTRUCT^TM^: 3.3 x 10^4^ ± 0.4 x 10^4^-fold, Matrigel^TM^: 2.8 x 10^4^ ± 0.4 x 10^4^-fold). Expression of the corresponding lineage-specific transcription factors *ARX* (α-cell lineage) and *HHEX* (δ-cell lineage) remained comparable between BIOINSTRUCT^TM^ and Matrigel^TM^ conditions. Consistent with these findings, SC-Islets generated on BIOINSTRUCT^TM^ contained GCG-positive α-like cells, CPEP-positive β-like cells, and SST-positive δ-like cells (Fig. 7d), demonstrating the formation of the major endocrine cell populations. To further assess the efficiency of pancreatic endocrine progenitor formation on BIOINSTRUCT^TM^, we quantified the frequency of PDX1- and NKX6.1-positive cells in spheroids four days after detachment (day 15 of differentiation) (Fig. 7e,f).

Both, BIOINSTRUCT^TM^ and Matrigel^TM^ cultures, exhibited high frequencies of PDX1-positive (91 ± 10% and 93 ± 5%), NKX6.1-positive (64 ± 20% and 61 ± 22%), and PDX1/NKX6.1 double-positive cells (63 ± 20% and 61 ± 22%), indicating efficient generation of pancreatic endocrine progenitors on BIOINSTRUCT^TM^-coated plates. Similarly, analysis of mature SC-Islets at day 48 revealed comparable frequencies of PDX1-positive (50 ± 22% and 47 ± 22%), NKX6.1-positive (42 ± 15% and 39 ± 19%), and PDX1/NKX6.1 double-positive populations (34 ± 15% and 30 ± 19%) between BIOINSTRUCT^TM^ and Matrigel^TM^, respectively (Fig. 7g,h).

Taken together, these data demonstrate that BIOINSTRUCT^TM^ coating supports the generation of hiPSC-derived pancreatic endocrine cells using established differentiation protocols and enables formation of SC-Islets comparable to Matrigel^TM^-based cultures.

## Discussion

Animal-derived basement membrane extracts (BMEs) represent the gold standard in stem cell culture. Numerous studies have successfully employed BMEs, particularly Matrigel^TM^, in a wide range of hiPSC-based applications. Their complex composition, which includes laminin, collagen IV, entactin, and bioactive growth factors^9,11^, enables their versatile use. However, this compositional complexity, is associated with considerable batch-to-batch variability, resulting in lot-dependent performance differences. Furthermore, the production of BMEs from mouse tumors^10^, as in the case of Matrigel^TM^ and Geltrex™, raises ethical concerns and limits their applicability for generating hiPSC derivatives intended for clinical use^12^. Recombinant proteins, such as laminin-511/521^13^ and vitronectin^16^, as well as laminin E8 fragments^14^, offer animal-free alternatives and represent a compromise between biological complexity and standardization. However, compared with BMEs, recombinant protein-based substrates provide a more limited range of integrin-binding options for hiPSCs, potentially reducing cell adhesion in certain cell lines and limiting their compatibility with specific lines. Moreover, recombinant protein production remains comparatively complex and costly at scale, requiring biological expression, purification, and stringent quality control, with associated risks of batch variability and process-related impurities.

Fully synthetic ECMs therefore offer an attractive alternative. In recent years, several approaches have been developed to establish synthetic alternatives to animal-derived BMEs for stem cell culture to circumvent the disadvantages of animal-based and recombinant surface coatings. These approaches have shown promising results in supporting the long-term culture of pluripotent stem cells (PSCs), but have not yet achieved widespread adoption for various reasons. Lambshead et al. and Klim et al. demonstrated the long-term culture of human embryonic stem cell (hESC) lines on synthetic culture substrates for 10 passages^35^ and 3 months^18^, respectively. However, their approaches relied on specific manufacturing processes that may impede scalability and commercial production. Other substrates, such as the synthetic coating developed by Shimizu et al., have been shown to support the culture of individual cell lines^20^, but evidence of their broader applicability remains limited.

The present study indicates that BIOINSTRUCT^TM^ represents a suitable synthetic alternative to animal-derived BMEs and recombinant proteins for the long-term maintenance of hiPSCs and their subsequent directed differentiation. Unlike some previously described synthetic substrates^18,35^, BIOINSTRUCT^TM^ does not require specialized coating procedures or cold chain logistics, which may facilitate its scalability and broader applicability. Collectively, our data demonstrate that BIOINSTRUCT^TM^ supports long-term hiPSC maintenance and the generation of specified cell types through directed differentiation across different germ layer lineages. However, whereas BMEs and recombinant protein-based substrates have been validated across a variety of cell lines, this study investigated BIOINSTRUCT^TM^ on a limited number of hiPSC lines. We show that BIOINSTRUCT^TM^ supports the maintenance of the IMR90-4 cell line and the directed differentiation of HIHCNi008-A, ACTN2-GFP, and UKKI-011-A cells. Furthermore, we collected evidence that BIOINSTRUCT^TM^ may also support the maintenance of the BIONi010B cell line. Although these findings demonstrate the suitability of BIOINSTRUCT^TM^ for the culture and differentiation of several cell lines and suggest its potential for broader applications, further validation using additional cell lines is required to establish its applicability across a wider range of hPSC lines.

hiPSCs cultured on BIOINSTRUCT^TM^ largely retained a stem cell-like morphology. Nevertheless, differences compared with cells cultured on Matrigel^TM^ were observed, including partially enlarged cells with a slightly lower nuclear-to-cytoplasmic ratio, the formation of broad pseudopodia, flattening of colony borders, and delayed detachment from the culture surface following Accutase treatment. Collectively, these observations suggest that BIOINSTRUCT^TM^ may promote stronger cell adhesion. Differences in substrate-cell interactions may arise from variations in the density of available ECM binding sites^36^. The density of available binding sites can influence mechanosignaling in cells^36^, which plays an important role in maintaining stemness and regulating differentiation in PSCs^37,38^. So far, we did not detect any changes in the maintenance of stemness or the retention of pluripotency in cells cultured on BIOINSTRUCT^TM^. In addition, cellular morphology has been shown to influence cell fate decisions during differentiation. For instance, Mamidi et al. have demonstrated that the endocrine commitment of pancreatic progenitors is strongly influenced by cell shape and the actin cytoskeleton^39^. In our study, pancreatic differentiation on BIOINSTRUCT^TM^ did not differ substantially from that on Matrigel^TM^, suggesting that the morphological differences observed during hiPSC maintenance do not markedly affect directed differentiation under the conditions investigated.

In addition to the morphological differences, a higher cell count at the end of each passage was observed on BIOINSTRUCT^TM^, resulting in a higher calculated population doubling rate compared with Matrigel^TM^. Increased proliferation represents one possible explanation for this observation. However, the higher cell count could also result from reduced cell death during culture or increased cell attachment following seeding. The latter possibility is supported by the differences in cell spreading observed on BIOINSTRUCT^TM^ and the delayed detachment of cells from the culture surface following Accutase treatment. As neither the number of attached cells following seeding nor the proliferative capacity of the cells was quantified, no definitive conclusions can be drawn regarding the effect of BIOINSTRUCT^TM^ on hiPSC proliferation in this study.

Retention of pluripotency of hiPSCs cultured on BIOINSTRUCT^TM^ was assessed by spontaneous differentiation. To evaluate multilineage differentiation in a standardized and quantitative manner and to investigate potential lineage biases associated with BIOINSTRUCT^TM^, we employed the germ layer scoring method introduced by D’Antonio et al.^25^ using an adapted modified primer panel. This method evaluates the expression of genes encoding proteins associated with early ectodermal, mesodermal, and endodermal development and calculates a germ layer score based on the average expression of genes assigned to each germ layer. Although this approach provides a relatively rapid and cost-effective means of assessing differentiation potential, it does not account for the pleiotropic functions of certain transcription factors. For example, *GATA4* is included as an endoderm-associated marker in the germ layer scoring method^25^, while it is also involved in the development of mesoderm-derived cardiac progenitor cells^40^. Although germ layer scoring may oversimplify the developmental roles of individual transcription factors, it nevertheless provides a standardized approach for identifying potential biases in early lineage differentiation associated with the culture substrate. Using this method, we observed a slightly reduced endodermal score in cells maintained on BIOINSTRUCT^TM^ for 10 passages, potentially indicating a shift in differentiation propensity or delayed onset of endodermal differentiation. Analyses of the naïve and primed pluripotency states of hiPSC cultured on BIOINSTRUCT^TM^, as well as directed differentiation toward endoderm-derived pancreatic endocrine progenitors, did not reveal any abnormalities in the pluripotency state or endodermal differentiation. We therefore conclude that BIOINSTRUCT^TM^ does not introduce a shift in lineage commitment during differentiation.

In conclusion, this study demonstrates the suitability of BIOINSTRUCT^TM^ as a xeno-free, synthetic surface coating for the maintenance culture of hiPSCs as well as their directed differentiation. Compared to the gold standard Matrigel^TM^, no negative differences in the maintenance of pluripotency and induction of multilineage differentiation have been detected. Nevertheless, to provide evidence for the general applicability to hiPSC culture, validation of BIOINSTRUCT^TM^ with a greater number of cell lines is required. At this point, BIOINSTRUCT^TM^ presents a promising alternative for the xeno-free culture of stem cells and their derivatives.

## Resource availability

### Lead contact

Requests for further information and resources should be directed to and will be fulfilled by the lead contact, Constantin Berger.

### Materials availability

This study did not generate new unique reagents.

### Data and code availability

All data reported in this paper will be shared by the lead contact upon request. Any additional information required to reanalyze the data reported in this paper is available from the lead contact upon request

## Acknowledgements / Funding

The authors thank Alevtina Höchner, Christa Albert and Evelin Gebert for their excellent technical support. J.G.W.S. and T.L.H. were supported by the British Heart Foundation (SP/F/22/150044). K.C. was supported by the UKRI BBSRC Norwich Research Park Bioscience Doctoral Training Program BB/T008717/1. J.K. gratefully acknowledges support from the SCIENTIA Fellowship Program, funded through the Bavarian Gender Equality Promotion (Bayerische Gleichstellungsförderung, BGF) of the Free State of Bavaria. Elina Nürnberg, Sarah Al Muthafer, and Sandra Segeletz are employed and funded by faCellitate GmbH. We thank the FlowCore facility of the Medical Faculty Mannheim at Heidelberg University, especially Iris Arantes de Castro, Stefanie Uhlig, Isabel Moskal, and Karen Bieback for their support with FACS analysis, and Kaan Cilek for his excellent technical support. The EBiSC Bank acknowledges Bioneer A/S as the source of the human induced pluripotent cell line BIONi010-B and KLINIKUM DER UNIVERSITAET ZU KOELN as the source of the human induced pluripotent cell line UKKi011-A, which were generated with support from the EBiSC project. The EBiSC has received support from the Innovative Medicines Initiative (IMI) Joint Undertaking (JU) under grant agreement n°115582 and from the IMI-2 JU under grant agreement No 821362, resources of which are composed of from the European Union’s Seventh Framework Programme (FP7/2007-2013), European Union’s Horizon 2020 research and innovation programme and EFPIA.

## Author contributions

CB: Conceptualization, Data curation, Formal analysis, Investigation, Methodology, Project administration, Resources, Visualization, Writing - Original Draft, Writing - Review C Editing; LW: Data curation, Formal analysis, Investigation, Visualization, Writing - Original Draft, Writing - Review C Editing; TH: Formal analysis, Investigation, Visualization, Writing - Original Draft, Writing-Review C Editing; FW: Formal analysis, Investigation, Writing - Original Draft; CS: Data curation, Formal analysis, Investigation, Visualization, Writing - Original Draft; LB: Data curation, Formal analysis, Investigation, Visualization, Writing - Original Draft; JS: Investigation; GB: Investigation; SL: Investigation; JK: Investigation; YS: Investigation; MK: Investigation; KC: Investigation; EN: Data curation, Formal analysis, Investigation, Methodology, Resources, Writing - Review C Editing; SAM: Data curation, Formal analysis, Investigation, Methodology, Resources, Writing - Review C Editing; SS: Conceptualization, Funding acquisition, Project administration; CN: Conceptualization, Methodology; JGWS: Conceptualization, Investigation, Funding acquisition, Methodology, Supervision, Writing - Review C Editing; SH: Conceptualization, Funding acquisition, Methodology, Supervision, Writing - Review C Editing; DZ: Conceptualization, Funding acquisition, Methodology, Supervision;

## Declaration of generative AI and AI-assisted technologies in the writing process

During the preparation of this work the authors used “GPT-5.6 Luna” in order to improve readability of the manuscript. After using this tool, the authors reviewed and edited the content as needed and take full responsibility for the content of the publication.

## Methods

### Key resources table

#### Antibodies

A list of antibodies with information about source and identifiers can be found in Supplemental Document 2.

#### Experimental models: Cell lines

| **Reagent or resource** | **Source** | **Identifier** |
|---|---|---|
| Human: IMR90-4 hiPSC (iPS(IMR90) clone #4) | WiCell | WISCi004-B |
| Human: UKKi011-A hiPSC | EBiSC | UKKi011-A |
| Human: BIONi010-B hiPSC | EBiSC / Sigma-Aldrich | BIONi010-B |
| Human: WTC-mEGFP-ACTN2-cl85 hiPSC | Allen Institute | AICS-0075-085 |
| Human: iPS-CO-22 2.1 hiPSC | Derived in-house; hPSCreg | HIHCNi008-A |

#### Experimental models: Organisms/strains

| **Reagent or resource** | **Source** | **Identifier** |
|---|---|---|
| Mouse: primary astrocytes from mixed cortical glial cultures (E18 embryos) | Isolated in-house (ref. 49) | N/A |

#### Chemicals, peptides, and recombinant proteins

| **Reagent or resource** | **Source** | **Identifier** |
|---|---|---|
| ***Coating and matrix*** |  |  |
| BIOINSTRUCT | faCellitate | N/A |
| Gelatine | Serva | 22151.02 |
| Laminin-521 | BioLamina | LN521-05 |
| Matrigel | Corning | 356231 |
| Poly-L-ornithine | Sigma-Aldrich | P8638 |
| ***Media, supplements, and buffers*** |  |  |
| 2-Mercaptoethanol (neuronal differentiation) | Sigma-Aldrich | M6250 |
| 2-Mercaptoethanol (spontaneous differentiation) | Gibco | 31350-010 |
| Anti-adherence rinsing solution | StemCell Technologies | 07010 |
| B27 minus insulin | Gibco | A1895601 |
| B27 minus vitamin A | Gibco | 12587010 |
| B27 supplement | Gibco | 17504044 |
| BrainPhys + SM1 | StemCell Technologies | 05792 |
| CMRL1066 | Gibco | 11530-037 |
| CryoStor CS10 | StemCell Technologies | 100-1061 |
| Dimethyl sulfoxide (DMSO) | Sigma-Aldrich | D8418 |
| DMEM/F12 with 15 mM HEPES | Gibco | 31330-038 |
| DMEM/F12 without glutamine | Gibco | 21331-020 |
| DMEM/F12 without HEPES | Gibco | 11320074 |
| Fetal calf serum (FCS) | Bio&Sell | FBS.S.0615 |
| GlutaMAX | Gibco | 35050-038 |
| HBSS | Biowest | L0607-500 |
| ITS | Gibco | 41400045 |
| ITS-X | Gibco | 51500056 |
| Knockout Serum Replacement | Gibco | 10828028 |
| Lipid concentrate | Gibco | 11905-031 |
| MCDB131 | Gibco | 10372-019 |
| MEM non-essential amino acids (neuronal differentiation) | Gibco | 11140050 |
| MEM non-essential amino acids (spontaneous differentiation) | Gibco | 11140035 |
| mTeSR Plus | StemCell Technologies | 100-0276 |
| mTeSR-1 | StemCell Technologies | 85850 |
| Neurobasal medium | Gibco | 21103049 |
| PBS with CaCl2 and MgCl2 | Gibco | 14040117 |
| PBS without CaCl2 and MgCl2 (for Flow cytometry and immunostaining) | Gibco | 14190144 |
| PBS without CaCl2 and MgCl2 (neuronal differentiation) | Sigma-Aldrich | D8537 |
| Penicillin/Streptomycin | Sigma-Aldrich | P4333 |
| RPMI 1640 | Gibco | 21875034 |
| RPMI minus glucose | Gibco | 11879020 |
| Trace Elements A | Corning | 25-021-CI |
| Trace Elements B | Corning | 25-022-CI |
| ***Dissociation reagents*** |  |  |
| Accutase (BIONi010-B passaging) | StemCell Technologies | 07920 |
| Accutase (IMR90-4 passaging, neuronal and pancreatic differentiation) | Sigma-Aldrich | A6964 |
| EDTA (flow cytometry buffer, BIONi010-B) | Applichem | A4892,0100 |
| EDTA (flow cytometry buffer, IMR90-4) | Applichem | A3553 |
| EDTA (iPS-CO-22 2.1 detachment) | Carl Roth | 8043.2 |
| Gentle Cell Dissociation Reagent | StemCell Technologies | 100-0485 |
| ***Small molecules, growth factors, and cytokines*** |  |  |
| Activin A (cardiac differentiation) | Proteintech | HZ-1138 |
| Activin A (pancreatic differentiation) | PeproTech | AF-120-14E |
| AGN (AGN 193109 sodium salt) | Santa Cruz Biotechnology | sc-210768 |
| ALK5 inhibitor II | Enzo Life Sciences | ALX-270-445 |
| Apo-transferrin | Sigma-Aldrich | T1147 |
| Ascorbic acid (cardiac differentiation) | Merck | A8960-5G |
| Ascorbic acid (pancreatic differentiation) | Sigma-Aldrich | A4544 |
| BDNF | PeproTech | 450-02 |
| Betacellulin | PeproTech | 100-50 |
| BMP4 | Proteintech | HZ-1045 |
| BSA (fatty acid carrier, culture) | Merck / Sigma-Aldrich | A7030-10G / A7030 |
| CHIR99021 (cardiac differentiation) | Cambridge Bioscience | BPS-27615-2 |
| CHIR-99021 (pancreatic differentiation) | Selleck | S1263 |
| D-(+)-Galactose | Merck | G5388-100G |
| DAPT | Sigma-Aldrich | D5942 |
| D-Glucose | Sigma-Aldrich | G7021 |
| EGF | PeproTech | AF-100-15 |
| FGF2 (cardiac differentiation) | Proteintech | HZ-1285 |
| FGF2 (neuronal differentiation) | PeproTech | 100-18B |
| FGF7 | PeproTech | AF-100-19 |
| GC1 | Tocris | 4554 |
| GDNF | PeproTech | 450-10 |
| Heparin (neuronal differentiation) | Sigma-Aldrich | H3393 |
| Heparin (pancreatic differentiation) | Sigma-Aldrich | H3149 |
| Insulin | Sigma-Aldrich | I6634 |
| IWR-1 | Merck | I0161 |
| Lactic acid | Merck | 71720-5G |
| L-Ascorbic acid 2-phosphate sesquimagnesium salt | Sigma-Aldrich | A8960 |
| LDN-193189 (neuronal differentiation) | Th. Geyer | SML0559 |
| LDN-193189 (pancreatic differentiation) | Selleck | 7507 |
| N-Acetylcysteine | Sigma-Aldrich | A9165 |
| NGF | PeproTech | 450-01 |
| Nicotinamide | Sigma-Aldrich | N0636 |
| Oleic acid | Merck | O1383-1G |
| Palmitic acid | Merck | P0500-10G |
| PD0325901 | Tocris | 4192 |
| Progesterone | Sigma-Aldrich | P8783 |
| Putrescine | Sigma-Aldrich | P5780 |
| Retinoic acid | Sigma-Aldrich | R2625 |
| SANT-1 | Sigma-Aldrich | S4572 |
| SB431542 | Sigma-Aldrich | S4317 |
| Sodium bicarbonate | Sigma-Aldrich | S5761 |
| Sodium pyruvate | Sigma-Aldrich | S8636 |
| Sodium selenite | Sigma-Aldrich | 214485 |
| T3 | Sigma-Aldrich | 6397 |
| TBP | Tocris | 5342 |
| TGF-β1 | PeproTech | 100-21-C-01M |
| Y-27632 | Tocris | 1254 |
| Y-27632 (BIONi010-B maintenance) | StemCell Technologies | 72304 |
| ZM-447439 | Sigma-Aldrich | 2051 |
| ZnSO4 | Sigma-Aldrich | Z0251 |
| γ-Secretase inhibitor XX | Sigma-Aldrich | 565789 |
| ***Fixation, staining, and flow cytometry reagents*** |  |  |
| BD Cytofix/Cytoperm | BD Biosciences | 51-2090KZ |
| BD Perm/Wash | BD Biosciences | 51-2091KZ |
| BSA (blocking) | Carl Roth | 8076.5 |
| DAKO mounting medium | N/A in text | N/A |
| DAPI | Invitrogen / Thermo Fisher | 62248 |
| FcR blocking reagent | Miltenyi Biotec | 130-059-901 |
| Fixable Viability Stain 510 | BD Biosciences | 564406 |
| Fluoromount-G with DAPI | Invitrogen | 00-4959-52 |
| Formaldehyde | Merck | 1004968350 |
| Histogel | Epredia | HG-4000-012 |
| Paraformaldehyde | Carl Roth | 0335.2 |
| Phalloidin-647 | Abcam | ab176759 |
| Qiazol lysis reagent | Qiagen | 79306 |
| RNase A | N/A in text | N/A |
| Roticlear Histofix 4% | Carl Roth | P087.3 |
| Saponin | Carl Roth | 4185.1 |
| Sodium azide | Sigma-Aldrich | SIAL71289-5G |
| Triton X-100 | Sigma-Aldrich | T8787 |
| Tween-20 | Sigma-Aldrich | P1379 |

#### Commercial Kits

| Reagent or resource | Source | Identifier |
|---|---|---|
| E.Z.N.A. MicroElute Total RNA Kit | Omega Bio-Tek | R6831 |
| iScript cDNA Synthesis Kit | Bio-Rad | 1708891 |
| RNeasy Plus Universal Mini Kit | Qiagen | 73404 |
| STEMdiff Cardiomyocyte Dissociation Kit | StemCell Technologies | 05025 |
| STEMdiff Trilineage Differentiation Kit | StemCell Technologies | 05230 |
| SYBR Green Ultramix | Bio-Rad | 1725274 |
| SYBR Select Master Mix | Applied Biosystems | 4472908 |
| UltraScript 2.0 cDNA Synthesis Kit | PCR Biosystems | PB30.11-02 |

#### Deposited data

| **Reagent or resource** | **Source** | **Identifier** |
|---|---|---|
| None stated in the methods |  |  |

#### Oligonucleotides

A list of primers with information about primer sequence and RefSeq identifier of the targeted transcript can be found Supplemental Document 2.

#### Software and algorithms

| **Reagent or resource** | **Source** | **Identifier** |
|---|---|---|
| FFMPEG | ffmpeg.org | RRID:SCR_016075 |
| FlowJo | BD | RRID:SCR_008520 |
| Gen5 software | BioTek | RRID:SCR_017317 |
| GraphPad Prism | GraphPad | RRID:SCR_002798 |
| ImageJ (v1.53f51) | NIH | RRID:SCR_003070 |
| Imaris v11.0.0 | Bitplane, Oxford Instruments | RRID:SCR_007370 |
| Multiwell-Analyzer v2.0.6.0 | Multi Channel Systems (MCS) | N/A |
| Multiwell-Screen v2.0.11.0 | Multi Channel Systems (MCS) | N/A |
| MUSCLEMOTION (ImageJ plugin) | Sala et al.<sup>32</sup> | N/A |

#### Other (consumables, devices)

| Reagent or resource | Source | Identifier |
|---|---|---|
| 24-well Multiwell-MEA plates, 12 electrodes/well | MCS | 24W700/100F-288 |
| AggreWell 400 plates | StemCell Technologies | 34415 |
| AggreWell 800 plates | StemCell Technologies | 34815 |
| Glass cover slips | Hartenstein | DK24 |
| Glass-bottom plates / 24-well plates | Ibidi | 82421 |
| Non-tissue-culture-treated plates | Corning | 351143 |
| Tissue-culture-treated plates | Thermo Fisher | 140675 |
| ***Devices*** |  |  |
| BD FACSCanto II flow cytometer | BD Biosciences | N/A |
| BZ-X810 fluorescence microscope | Keyence | N/A |
| Cytation 7 imager | BioTek | BTCYT7U |
| MACSQuant 10 flow cytometer | Miltenyi Biotec | N/A |
| Multiwell-MEA-System (with headstage) | Multi Channel Systems (MCS) | N/A |
| NanoDrop One | Thermo Fisher Scientific | ND-ONE-W |
| Nikon Ti2e-SORA with Yokogawa W1 spinning disc unit (40x water immersion objective) | Nikon / Yokogawa | N/A |
| Orbital shaker (100 rpm) | N/A in text | N/A |
| QuantStudio 6 Pro (qPCR, neuronal RNA) | Applied Biosystems | N/A |
| QuantStudio 7 (qPCR) | Applied Biosystems | 4485701 (cardiac RNA);<br>N/A for other uses |

### Experimental model and study participant details

#### Human iPSC lines

IMR90-4 (iPS(IMR90) clone #4; WISCi004-B) cells were obtained from WiCell and originates from a female donor with no reported disease. Working stocks were cryopreserved at passage 12 and experiments conducted at passage 14 – 28.

UKKi011-A cells were obtained from EBiSC and originates from a male Caucasian donor with no reported disease. Working stocks were cryopreserved at passage 13 and experiments conducted at passage 18 – 22.

BIONi010-B cells were obtained from EBiSC/Sigma-Aldrich and originates from a healthy 18-year-old male donor of African ethnicity with no reported disease. Working stocks were cryopreserved at passage 29 and experiments were conducted within 10 passages after thawing.

WTC-mEGFP-ACTN2-cl85 (AICS-0075-085) cell were obtained from the Allen Institute. It is derived from the parental WTC line (Coriell GM25256; leg skin fibroblasts of a 30-year-old male Japanese donor) and carries a monoallelic C-terminal mEGFP tag on ACTN2 introduced by CRISPR/Cas9. No disease mutation is known. Working stocks were cryopreserved at passage 35 and experiments conducted after thawing.

iPS-CO-22 2.1 (hPSCreg: HIHCNi008-A) cells were derived from a skin biopsy of a Caucasian male donor without known disease. Fibroblasts were cultured to passage 4 and reprogrammed episomally by electroporation. Working stocks were cryopreserved at passage 8 and experiments conducted at passage 10 – 15.

#### hiPSC culture and quality control

IMR90-4 cells were maintained on Matrigel^TM^- or BIOINSTRUCT^TM^-coated plates in mTeSR-1 medium. IMR90-4 cells were passaged as single cells (Accutase) and seeded at a density of 5.21 x 10^3^ cells/cm^2^ in mTeSR-1 medium supplemented with 10 µM Y-27632 for 24 h. IMR90-4 cells were cryopreserved in DMEM without Glutamine containing 20% Knockout Serum Replacement and 10% DMSO. For thawing, IMR90-4 cells were plated in mTeSR-1 medium containing 10 µM Y-27632. IMR90-4 cells were cultured without antibiotics. Sterility was monitored daily by microscopy. Cultures were tested for mycoplasma contamination every three months by mycoplasma PCR. Cell line identity was not tested. Genomic integrity was assessed by G-banding karyotyping (20 metaphases, 300 – 500 bands) at passage 10 after thawing of the working stock. The undifferentiated state was assessed by gene expression (*OCT4*, *SOX2*, *NANOG*) and FACS (NANOG, TRA1-81) analysis at passage 5 and 10 after thawing of the working stock. Pluripotency was assessed by spontaneous differentiation at passage 11 after thawing of the working stock.

UKKI011-A cells were maintained on Matrigel^TM^-coated plates in mTeSR-1 medium and passaged using Gentle Cell dissociation reagent as clumps at a split ratio of 1:6 – 1:20 without ROCK inhibitor. UKKI011-A cells were cryopreserved in DMEM without Glutamine containing 20% Knockout Serum Replacement and 10% DMSO. For thawing, UKKI011-A cells were plated in mTeSR-1 medium containing 10 µM Y-27632. UKKI011-A cells were cultured without antibiotics. Sterility was monitored daily by microscopy. Cultures were tested for mycoplasma contamination every three months by mycoplasma PCR. Cell line identity and genomic integrity were not tested after purchase. The undifferentiated state was assessed by gene expression (*OCT4*, *SOX2*, *NANOG*) and FACS analysis (NANOG, TRA1-81) before experimental use.

BIONi010-B cells were maintained on Matrigel^TM^- or BIOINSTRUCT^TM^-coated plates in mTeSR Plus medium. BIONi010-B cells were passaged as single cells (Accutase) and plated at a density of 1.04 x 10^4^ cells/cm^2^ in mTeSR Plus medium supplemented with 10 µM Y-27632 for 24 h. BIONi010-B cells were cryopreserved with CryoStor CS10. For thawing, BIONi010-B cells were plated in mTeSR Plus containing 10 µM Y-27632. Cell line identity was confirmed by short tandem repeat (STR) testing. Sterility was monitored daily by microscopy. Cultures were tested for mycoplasma contamination using an external service from StemGenomics. Genomic integrity was assessed by G-banding karyotyping (20 metaphases, 300 – 500 bands) at passage 10 after thawing of the working stock. The undifferentiated state was assessed by FACS analysis (OCT4, SOX2, SSEA-4) at passage 10 after thawing of the working stock. Pluripotency was confirmed by directed trilineage differentiation at passage 10 after thawing of the working stock.

WTC-mEGFP-ACTN2-cl85 cells were maintained on Matrigel^TM^-coated plates in E8 medium. WTC- mEGFP-ACTN2-cl85 cells were passaged as single cells (Accutase) and plated at a cell density of 2 x 10^4^ cells/cm^2^ in E8 medium. WTC-mEGFP-ACTN2-cl85 cells were cryopreserved in E8 medium containing 10% DMSO. For thawing, WTC-mEGFP-ACTN2-cl85 cells were plated in E8 medium containing 10 µM Y-27632. WTC-mEGFP-ACTN2-cl85 cells were cultured without antibiotics. Sterility was monitored daily by microscopy. Cultures were tested for mycoplasma contamination every three months by IDEXX qPCR. Cell line identity was confirmed by short tandem repeat (STR) testing. Genomic integrity was assessed by G-banding karyotyping. The undifferentiated state was assessed by FACS analysis (OCT4, SOX2, NANOG). Genetic modification (integration of monoallelic C-terminal mEGFP tag on ACTN2) was confirmed by PCR and immunocytochemistry.

iPS-CO-22 2.1 cells were maintained on Matrigel^TM^- or BIOINSTRUCT^TM^-coated plates in in-house prepared E8 medium. iPS-CO-22 2.1 cells were passaged using PBS/EDTA at a split ratio of 1:10 – 1:20 in in-house prepared E8 medium supplemented with 10 µM Y-27632 for 24 h. iPS-CO-22 2.1 cells were cryopreserved in medium containing Knockout Serum replacement and 10% DMSO. For thawing, iPS-CO-22 2.1 cells were plated in E8 medium containing 10 µM Y-27632. iPS-CO-22 2.1 cells cultured without antibiotics. Sterility was monitored daily by microscopy. Cultures were tested for mycoplasma contamination every four months. Cell line identity was not tested. Genomic integrity was assessed at prepration of the working stock (passage 8) by SNP (Illumina Bead Array Human Omni2.4 Exome-8 Bead Chip v1.4) using an external service by LIFE C BRAIN GmbH. The undifferentiated state was assessed by immunocytochemistry (OCT4, TRA1-81) and gene expression analysis of seven pluripotency markers.

### Method details

#### Coating of plates

For Matrigel^TM^-coating, tissue-culture-treated plates, glass bottom plates or glass cover slips were covered with Matrigel^TM^ diluted in DMEM/F12 without Glutamine at a ratio of 83.33 ng/mL for 1h at RT. BIOINSTRUCT^TM^-coated cell culture surfaces were generated by incubating non-tissue-culture treated plates, glass bottom plates or glass cover slips with BIOINSTRUCT^TM^ solution (faCellitate) for 5 -60 min at RT. For Gelatine-coating, tissue cultured plates or glass-bottom plates were treated with a 0.1% Gelatine solution (w/v) for 1 h at 37°C. For laminin coating, laminin stock solution was diluted to a final concentration of 5 µg/mL in Phosphate Buffered Solution (PBS) with CaCl_2_ and MgCl_2_. Laminin-coated cell culture surfaces were generated by incubating plates for 2 h at 37 °C.

#### Maintenance culture on Matrigel- and BIOINSTRUCT-coated culture plates

IMR 90-4 hiPSCs were cultured in mTeSR-1 medium on Matrigel^TM^ or BIOINSTRUCT^TM^-coated plates at 37°C, 5% CO_2_, 95% humidity. hiPSCs were passaged every 4^th^ day as single cells. Therefore, hiPSCs on Matrigel^TM^ or BIOINSTRUCT^TM^ were incubated with Accutase for 5 or 7 min at 37 °C, respectively. Cells on BIOINSTRUCT^TM^ required additional mechanical detachment using a cell scraper. Following resuspension in mTeSR-1 medium 5 x 10^4^ cells per 6-well (≙ 5.21 x 10^3^ cells/cm^2^) were plated in mTeSR-1 medium supplemented with 10 µM Y-27632.

BIONi010B hiPSCs were cultured in mTeSR-Plus medium on Matrigel^TM^ or BIOINSTRUCT^TM^-coated plates at 37 °C and 5% CO2. hiPSCs were passaged every 4th to 5th day as single cells. Therefore, hiPSCs on Matrigel^TM^ or BIOINSTRUCT^TM^ were incubated with Accutase for 7 or 10 min at 37 °C, respectively. Following resuspension in mTeSR-Plus medium 10^5^ cells per 6-well (≙ 1.04 x 10^4^ cells/cm^2^) were plated in mTeSR-Plus medium supplemented with 10 µM Y-27632.

#### Cryopreservation and thawing of IMRS0-4 cells

For cryopreservation, hiPSC were harvested as single cells using Accutase. Cells were counted and frozen at -80 °C at a final concentration of 2.5 x 10^5^ cells/mL in mTeSR-1 medium containing 10% Dimethyl Sulfoxide using freezing containers. After 24 h storage at -80 °C, hiPSCs were moved to liquid nitrogen. For reviving, cryotubes were thawed in a 37 °C waterbath and 1.25 x 10^5^ cells /6 well (≙ 1.3 x 10^4^ cells/cm^2^) were plated in 2 mL mTeSR-1 medium supplemented with 10 µM Y-27632 on Matrigel^TM^ or BIOINSTRUCT^TM^-coated plates.

#### Cell growth characterization

To determine cell growth characteristics, a defined number of hiPSCs were seeded and counted at the end of each passage (every 4 days), respectively. Doubling rate (i) and doubling time (ii) were calculated using the following formula:

i. 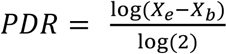
ii. 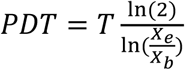

where *PDR* is the Population doubling rate, *PDT* is the Population doubling time, *Xe* is the cell count at the end of the passage, *Xb* is the cell count at the beginning of the passage and *T* the duration of passage in days. The cumulative doubling rate was calculated as the sum of the doubling rates of all previous passages.

#### Spontaneous differentiation

Spontaneous differentiation was performed as described previously^1,41^ with slight modifications. hiPSCs cultured on Matrigel^TM^ or BIOINSTRUCT^TM^ were harvested as single cells using Accutase. 9 x 10^5^ cells in 2 mL mTeSR-1 medium supplemented with 10 µM Y-27632 were transferred to a 24-well of a Aggrewell 800 plate for EB formation (≙ 3,000 cells/spheroid). After two days, formed EBs were transferred to petridishes and cultured in spontaneous differentiation medium, consisting of DMEM/F12 without Glutamine supplemented with 1x Glutamax, 20% Knockout Serum replacement, 1% Penicillin/Streptomycin, 100 mM non-essential amino acids, and 0.1 mM 2-Mercaptoethanol in suspension without agitation for 4 days. At day 6, EBs were collected and transferred to plates coated with Matrigel^TM^, BIOINSTRUCT^TM^ or 0.1 % Gelatine. EBs were cultured for up to 20 days with medium change every other day. To prevent detachment of loosely attached spheroids, only half medium changes were performed within the first 6 days.

#### Trilineage Differentiation

Trilineage differentiation of BIONi010B cells was carried out using the STEMdiff™ Trilineage Differentiation Kit according to the manufacturer’s instructions. Briefly, 24 well plates were coated with BIOINSTRUCT^TM^ or laminin as previously described. Subsequently, cells were seeded as single cells in mTeSR-Plus supplemented with 10 µM Y-27632, at cell densities recommended by the manufacturer. On the next day, medium was changed to the respective germ line-specific medium to initiate differentiation. Medium change was carried out daily for 5 days for mesoderm and endoderm differentiation. Differentiation into ectoderm was carried out for seven days.

##### Directed neuronal differentiation

Cortical layer V/VI neurons were generated from hiPSCs using a protocol adapted from Shi et al.^28^ and Rehbach et al.^30^. In brief, hiPSCs were dissociated using 0.02 % EDTA in PBS and seeded at a density of 3×10^5^ cells/cm² in Essential 8 medium (1x DMEM/F12 with 15 mM HEPES, 1× insulin-transferrin-selenium, 64 µg/mL L-ascorbic acid 2-phosphate sesquimagnesium salt hydrate, 2 ng/mL TGF-β1, 10 ng/mL FGF2, and 0.1 µg/mL heparin supplemented with 10 µM Y-27632. The following day, the medium was replaced with 3N medium, consisting of 50% N2 medium (see below), 50% Neurobasal medium, 0.5 x B27 supplement, 1x GlutaMax, 0.5 x non-essential amino acids, 50 µM 2-mercaptoethanol, 1 x Penicillin/Streptomycin and 1.6 mg/mL glucose. N2 medium consisted of DMEM/F12 without HEPES supplemented with 0.1 mg/mL apo-transferrin, 0.025 mg/mL insulin, 2 µM progesterone, 0.1 mM putrescine, and 0.03 µM sodium selenite. For neural induction, 3N medium was supplemented with 500 nM LDN-193189 and 10 µM SB431542 and replaced daily. On day 8, 20 ng/mL fibroblast growth factor 2 was added to the induction medium.

On day 9, cultures were dissociated using Accutase (10 min) and replated onto poly-L-ornithine-coated six-well plates additionally coated with either Matrigel^TM^ or BIOINSTRUCT^TM^. Cells were maintained in 3N medium supplemented with 20 ng/mL FGF2 and 10 µM Y-27632. The following day, the medium was replaced with 3N medium containing 20 ng/mL FGF2. From day 11 onwards, cells were maintained in 3N medium supplemented with100 ng/mL heparin, with medium changes every 48 h. Neural precursor cells were dissociated and cryopreserved on day 19.

For neuronal maturation, cryopreserved neural precursors were thawed and seeded in 3N medium supplemented with 10 µM Y-27632, with medium changes performed every 48 h. On day 26, cells were dissociated using Accutase and replated at assay-specific densities onto poly-L-ornithine-coated wells additionally coated with either Matrigel^TM^ or BIOINSTRUCT^TM^. Cells were seeded at 1×10^5^ cells/cm² for immunocytochemistry and at 3×10^5^ cells/cm² for RNA isolation. On days 27 and 29, the medium was replaced with 3N medium supplemented with 10 µM PD0325901 and 10 µM DAPT. From day 31 until fixation/sampling on day 36, cells were maintained in 3N medium with medium changes every 48 h.

#### Directed cardiac differentiation

Ventricular cardiomyocytes (hiPSC-CMs) were generated from the male ACTN2-GFP-hiPSC line (AICS-0075-085 purchased from the Allen Institute) using methods adapted from Dark et al.^31^. In brief, hiPSCs were seeded onto Matrigel^TM^ or BIOINSTRUCT^TM^-coated plates at a density of 2 x 10^4^ cells/cm^2^. The cells were grown to 70% confluency, at which point the differentiation procedure was commenced (day 0). The medium was replaced with cardiomyocyte differentiation medium comprised of RPMI supplemented with 1 x B27 minus insulin, 100 nM AGN Sodium Salt, 2 µM CHIR-99021, 5 ng/mL BMP4, 5 ng/mL Activin A and 5 ng/mL FGF2. After 24 hours, the medium was replaced with RPMI supplemented with 100 nM AGN and 1x B27 minus insulin (RPMI/B27-I). The following day, the medium was replaced with RPMI/B27-I supplemented with 2 µM IWR-1 and 65 µg/mL ascorbic acid. Subsequently, the medium was replaced with fresh RPMI/B27-I plus 65 µg/mL ascorbic acid every 48 hours until day 8. On day 8 and day 14, the medium was replaced with RPMI supplemented with 65 µg/mL ascorbic acid, 100 nM AGN, and 1 x B27 minus vitamin A (RPMI/B27-A). On day 10 and day 12, the medium was changed for RPMI minus glucose supplemented with 100 nM AGN, 1x B27 minus vitamin A, 65 µg/mL ascorbic acid and 4 mM lactic acid. On day 15, the cells were dissociated sing the STEMdiff™ Cardiomyocyte Dissociation Kit according to manufacturer’s instructions and replated on Matrigel^TM^ or BIOINSTRUCT^TM^-coated plates. After 24 hours following dissociation, the medium was changed to RPMI supplemented with 1 x B27. On day 20, the medium was changed to Galactose and Fatty Acid Media (GFAM), which is comprised of RPMI minus glucose, supplemented with 10 mM D-(+)- Galactose, 100 µM oleic acid and 50 µM palmitic acid. Before supplementation, fatty acids were dissolved in a BSA carrier (7.5% in dPBS). The medium was replaced with fresh GFAM every 48 hours from Day 20 onwards.

#### Directed pancreatic differentiation

Directed pancreatic differentiation was performed according to the protocols published by Barsby et al. and Balboa et al.^33,34^ using the UKKi011-A hiPSC line. For this, hiPSCs were collected as single cells using Accutase and 1.5 x 10^6^ cells seeded in 3 mL mTeSR-1 medium supplemented with 10 µM Y-27632 on Matrigel^TM^ or BIOINSTRUCT^TM^-coated plates (≙ 1.56 x 10^5^ cells/cm^2^). 24 h after seeding, the medium was changed to Basal medium (BM) 1 containing 100 ng/mL Activin A and 3 µM CHIR-99021. At the subsequent days the following media were used: Day 2: BM1 supplemented with 100 ng/mL Activin A, 0.3 µM CHIR-99021; day 3: BM1 supplemented with 100 ng/mL Activin A; day 4-6: BM1 supplemented with 0.25 mM Ascorbic Acid, 50 ng/mL FGF7; day 7-8: BM2 supplemented with 0.25 mM Ascorbic Acid, 50 ng/mL FGF7, 250 nM SANT-1, 1 µM Retinoic Acid, 100 nM LDN-193189, 200 nM TBP; day 9-12: BM2 supplemented with 0.25 mM Ascorbic Acid, 2 ng/mL FGF7, 100 ng/mL EGF, 10 ng/mL Activin A, 10 µM Y-27632, 10 mM Nicotinamide, 250 nM SANT-1, 0.1 µM Retinoic Acid, 200 nM LDN-193189, 100 nM TBP; day 13-16: BM3 supplemented with 20 ng/mL Betacellulin, 10 µM Y-27632, 250 nM SANT-1, 50 nM Retinoic Acid, 100 nM LDN-193189, 1 µM GC1, 100 nM γ-secretase inhibitor XX, 10 µM ALK5 inhibitor II; day 17-23: BM3 supplemented with 100 nM LDN-193189, 1 µM GC1, 100 nM γ-secretase inhibitor XX, 10 µM ALK5 inhibitor II; day 24-48: BM4 supplemented with 1 mM N-Acetylcystein, 10 nM T3, 0.5 µM ZM-447439. The basal media were composed as follows: BM1: MCDB131 medium supplemented with 1x Glutamax, 10 mM Glucose, 1.5 g/L NaHCO_3_, 0.5% BSA; BM2: MCDB131 medium supplemented with 1x Glutamax, 10 mM Glucose, 2.5 g/L NaHCO_3_, 2% BSA, 0.5% ITS-X; BM3: MCDB131 medium supplemented with 1x Glutamax, 20 mM Glucose, 1.5 g/L NaHCO_3_, 2% BSA, 0.5% ITS-X, 10 µM ZnSO_4_, 10 µg/mL Heparin sodium salt, 1x 1% Penicillin/Streptomycin; BM4: CMRL1066 medium supplemented with 1x Glutamax, 1% Penicillin/Streptomycin, 2% BSA, 0.5% ITS-X, 10 µM ZnSO_4_, 10 µg/mL Heparin, 0.5 mM Sodium Pyruvate, 1x Lipid concentrate, 1x Trace Elements A, 1x Trace Elements B. At day 11, cells were detached by incubation with Accutase for 15 min and transferred to 24-well Aggrewell 400 plates at a density of 1.2 x 10^6^ cells/2 mL/well. Formed spheroids were transferred to 6-well plates pre-treated with anti-adherence solution and cultured on an orbital shaker at 100 rpm.

#### Gene expression analysis

For hiPSCs and hiPSC-derived pancreatic islet-like clusters, total RNA was isolated using the E.Z.N.A.® MicroElute® Total RNA Kit according to the manufacturer’s guideline. 500 - 800 ng RNA were transcribed to cDNA with the iScript cDNA Synthesis Kit. qPCR was performed in duplicates with a QuantStudio 7 and SYBR-Green Ultramix using a two-step protocol with an annealing and elongation temperature of 60°C. Relative gene expression was calculated using the ddCT-method with *RPL4*, *RPLc*, and *RPS2S* for normalization. The germ layer score was determined according to D’antonio et al.^25^ with slight modifications.

For hiPSC-CMs, cells were lysed with Qiazol lysis buffer and RNA was isolated using the RNeasy Plus Universal Mini Kit according to manufacturer’s instructions. RNA was quantified using the Nanodrop One. Reverse transcription was performed on 1 µg RNA per sample using the Ultrascript 2.0 cDNA Synthesis Kit and the resulting cDNA was diluted 1:6 in nuclease-free water. Quantitative real-time PCR analysis was performed using a QuantStudio 7 using a two-step protocol with an annealing temperature and elongation temperature of 60°C. All real-time PCR reactions were performed in triplicate, normalized to housekeeper gene *GAPDH* and assessed using a comparative ddCt method.

For hiPSC-derived neuronal cells, gene expression analysis was performed by RT-qPCR using gene-specific primer pairs targeting neuronal, glial and housekeeping transcripts with SYBR select master mix on the Quant Studio 6 Pro System. Data was analyzed using the ddCt method.

Primer sequences were either custom-designed or retrieved from published studies^25,42–48^. A list of primers can be found in the Supplemental Document 2.

#### Immunoffuorescent staining

hiPSCs on coverslips or spontaneously differentiated cells on glass bottom plates were fixated by incubation in 4% Roticlear Histofix for 20 min. Trilineage differentiated hiPSCs were fixed using 4% PFA diluted in PBS without CaCl_2_ and MgCl_2_ (PBS^-^) for 30 min at 37 °C. For staining, samples were washed with PBS^-^ containing 0.5% Tween-20 (PBST) and subsequently treated with 0.2% Triton X-100 diluted in PBS^-^ for 5 min. After blocking of unspecific binding sites by incubation in 5% BSA dissolved in PBS^-^ for 30 min, samples were incubated with primary antibodies dissolved in blocking solution overnight at 4 °C. The next day, samples were washed three times for 5 min with PBST and treated with secondary antibodies diluted in antibody dilution solution for 2 h. For fluorescent labelling of Actin cytoskeleton, cells were incubated with Phalloidin-647 diluted 1:5000 in parallel to secondary antibody staining. Samples were washed three times for 5 min with PBST. hiPSCs on coverslips were mounted in Fluoromount G containing DAPI. Spontaneously differentiated cells and trilineage were stained with DAPI diluted 1:1000 in PBS-. Stained samples were imaged with a BZ-X810 Keyence fluorescence microscope.

hiPSC-derived islet-like clusters were fixated in Histofix for 45 min, mounted in Histogel, dehydrated in an ascending row of ethanol and embedded in paraffin. 3 µm paraffin sections were prepared, dewaxed and rehydrated and subsequently boiled in pH 6.0 citrate buffer at 95°C for 15 min for antigen retrieval. Sections were stained and imaged as described above.

For hiPSC-CMs, cells dissociated as previously described and seeded at a density of 10^4^ cells per cm^2^ onto 96-well plates coated with Matrigel^TM^ or BIOINSTRUCT^TM^ and cultured to day 30. On day 30, cells were washed once with PBS^-^ and then fixed with 4% formaldehyde for 15 min at room temperature. The cells were washed three times with PBS^-^ and incubated with 1 µg/mL, DAPI for 10 min at room temperature. The cells were washed three times with PBS^-^ and left under PBS^-^ during imaging. Cells were imaged using the Cytation 7 imager and analyzed using Gen 5 Software.

For hiPSC-derived neurons, cells were fixed using 4% paraformaldehyde for 10 min at 37°C, permeabilized and blocked with 5% BSA in PBS^-^ containing 0.1% Triton X-100 (PBS-X). Primary antibodies were diluted in the same blocking solution and incubated overnight at 4°C. After washing steps, matching secondary antibodies were applied in PBS-X in a dilution of 1:1000 for 45 min at RT followed by 15 min of DAPI/RNAseA treatment (1 µg/mL and 0.04 mg/mL respectively) for counterstaining. Cover glasses were mounted using DAKO mounting medium and allowed to dry overnight. Samples were imaged using the Nikon Ti2e-SORA system equipped with the Yokogawa W1 spinning disc unit and 40x water immersion objective. For characterization, selected fields of view were imaged in 25 µm z-stacks followed by subsequent maximum intensity projection.

Antibodies and dilutions are listed in Supplemental Document 2.

#### Image processing and quantification

Presented images were post-processed with Image J (v1.53f51) using the following commands: change to 8-bit/RGB, assign LUT, adjust brightness/contrast, subtract background (rolling ball), merge channels, stack to RGB.

The automated cell count of spheroids was conducted on unprocessed images. Initially, spheroids from images containing several spheroids were separated using the ImageJ cropping function in order to analyze individual spheroids. Quantification of immunofluorescent images of pancreatic spheroids was performed using Imaris (v11.0.0). DAPI-stained nuclei were segmented using the Imaris Spots detection module, with background subtraction enabled. An estimated nuclear diameter of 12 µm and 16 µm was used for day 15 (d15) and day 48 (d48) spheroids, respectively, to account for differences in cell size between timepoints. Intensity and quality thresholds for spot detection were determined visually on representative images and subsequently applied to all images using batch processing. Detected nuclei were classified as positive or negative for the nuclear markers NKX6.1 and PDX1 based on manually defined intensity threshold for the respective fluorescent channel. Separate thresholds were determined for d15 and d48 spheroids and applied to all images of the respective staining using batch processing. Segmentation results were visually reviewed to confirm plausibility. The number of positive cells was determined for each spheroid and marker. To assess co-expression of markers, cells were counted as negative, single-positive, or double-positive based on the combination of classification results.

Image analysis of hiPSC-CMs was performed on unprocessed images using the Cytation 7 Gen 5 Software. hiPSC-CMs were analyzed for the number of ACTN2-positive cells, mean fluorescence of ACTN2 staining, nucleus size and nucleus circularity.

#### Flow cytometry

IMR 90-4 cells were detached as single cells using Accutase and washed twice with Flow cytometry buffer (FC buffer), consisting of 1% FCS and 2 mM EDTA diluted in PBS^-^. hiPSCs stained for intracellular markers were incubated in BD Cytofix/Cytoperm for 20 min on ice. hiPSCs stained for extracellular markers were incubated in FcR-blocking reagent diluted 1:80 in FC buffer for 15 min on ice. Cells were washed with either BD Perm/wash for intracellular or FC buffer for extracellular staining. 1 – 2 x 10^5^ cells were transferred to FACS tubes and incubated with antibodies diluted in FC buffer for 30 min on ice. Following three times washing with FC buffer, cells were resuspended in FC buffer. Flow cytometry measurements were performed using a MACSQuant 10. All centrifugation steps were carried out at 350 x g for 3 min.

For BIONi010B cells, hiPSC were dissociated into single cells using Accutase for 7 – 10 min at 37 °C and resuspended in mTeSR-Plus with 10 µM Y-27632. Cells were centrifuged for 3 min at 280 x g, followed by resuspension of the cell pellet in cold PBS^-^. Viable cells were quantified by trypan blue exclusion using a Neubauer chamber and adjusted to 1 × 10^6^ cells per mL in PBS^-^. Subsequently, cells were stained with Fixable Viability Stain 510 for 15 min at room temperature in the dark, followed by washing in FACS buffer consisting of Hanks Balanced Salt Solution supplemented with 2% BSA, 2 mM EDTA, and 0.02% sodium azide. Cells were fixed in 2% PFA for 10 min and permeabilized in PBS^-^ containing 0.1% saponin and 1% BSA for 15 min at room temperature. 1 – 2 x 10^5^ cells were transferred to FACS tubes and incubated with antibodies diluted in FACS buffer for 60 min. Following antibody incubation, cells were washed twice using FACS buffer and analyzed on a BD FACSCanto™ II flow cytometer. All centrifugation steps during the staining procedure were performed at 400 × g for 3 min. Unless otherwise specified, samples were handled on ice and protected from light.

Data analysis was performed using FlowJo. A list of antibodies and the corresponding dilutions is given in Supplemental Document 2.

#### Karyotyping

Karyotyping was conducted by stemgenomics (Grabels, France). To this purpose, hiPSCs cultured for 10 passages on Matrigel^TM^, laminin, or BIOINSTRUCT^TM^ were shipped at 40% - 60 % confluency in mTeSR-1 or mTeSR-Plus medium. Karyotyping was performed using G-banding staining of 20 metaphases with a chromosome resolution of 300 – 500 base pairs.

#### Cardiomyocyte contraction analysis

To measure contraction, day 30 hiPSCs-CMs were recorded at 60 frames per second for 10 – 20 seconds. For each experiment, 5 wells were recorded per condition (BIOINSTRUCT^TM^ vs Matrigel^TM^), and the results averaged. The experiment was repeated 6 times. The recordings were converted from mp4 to tiffs using FFMPEG (http://ffmpeg.org/) and ImageJ, and then analyzed using the ImageJ plugin MUSCLEMOTION^32^ using default settings. Results were processed in Excel and GraphPad Prism.

#### Multielectrode array recordings

Spontaneous network activity was recorded using 24-well Multiwell-MEA plates with 12 electrodes per well. Plates were hydrated overnight in PBS at 37 °C and 5% CO₂ before coating. Wells were coated with poly-L-ornithine and either Matrigel ^TM^ or BIOINSTRUCT^TM^ using 50 µL per well, with six wells per condition. Matrigel ^TM^ was applied for 1 h at 37 °C and BIOINSTRUCT^TM^ for 5 min at room temperature. Before seeding, coating solutions were aspirated and wells were washed once with 3N medium.

On day 26 of differentiation, neurons were dissociated with Accutase for 5 min at 37 °C and seeded at 6 x 10^4^ cells per well in 50 µL 3N medium containing 10 µM Y-27632. After 30 min, 3 x 10^4^ primary mouse astrocytes, obtained from mixed cortical glial cultures of E18 embryos^49^, were added per well. After a further 30 min, the volume was adjusted to 500 µL. Cultures were maintained in 3N medium supplemented with 10 ng/mL each of BDNF, GDNF, and NGF. After 1 week, the medium was gradually replaced with BrainPhys supplemented with SM1 and the same neurotrophic factors. Half-medium changes were performed three times per week after recording.

Recordings were performed at 37 °C using a Multiwell-MEA-System and Multiwell-Screen software version 2.0.11.0. Plates were equilibrated on the headstage for 10 min before spontaneous activity was recorded for 5 min at 20 kHz. Recordings were performed two to three times per week from day 26 until day 87.

Signals were filtered using a first-order high-pass filter at 10 Hz and a second-order low-pass filter at 3500 Hz. Spikes were detected in Multiwell-Analyzer (version 2.0.6.0) with an electrode-specific threshold of 5x the standard deviation of the baseline noise. Bursts were identified on single electrodes with the maximum interval method (at least 4 spikes, maximum interspike interval of 50 ms). Network bursts were defined as synchronous bursts on at least six electrodes per well.

## Supplemental information

Document S1. Supplemental Figures 1-4

Document S2. Primer sequence and antibody information

