## Supplemental Document 1 for "A synthetic xeno-free extracellular matrix surrogate supports robust human induced pluripotent stem cell maintenance and multilineage differentiation"

### Document S1. Supplementary Figures 1-4

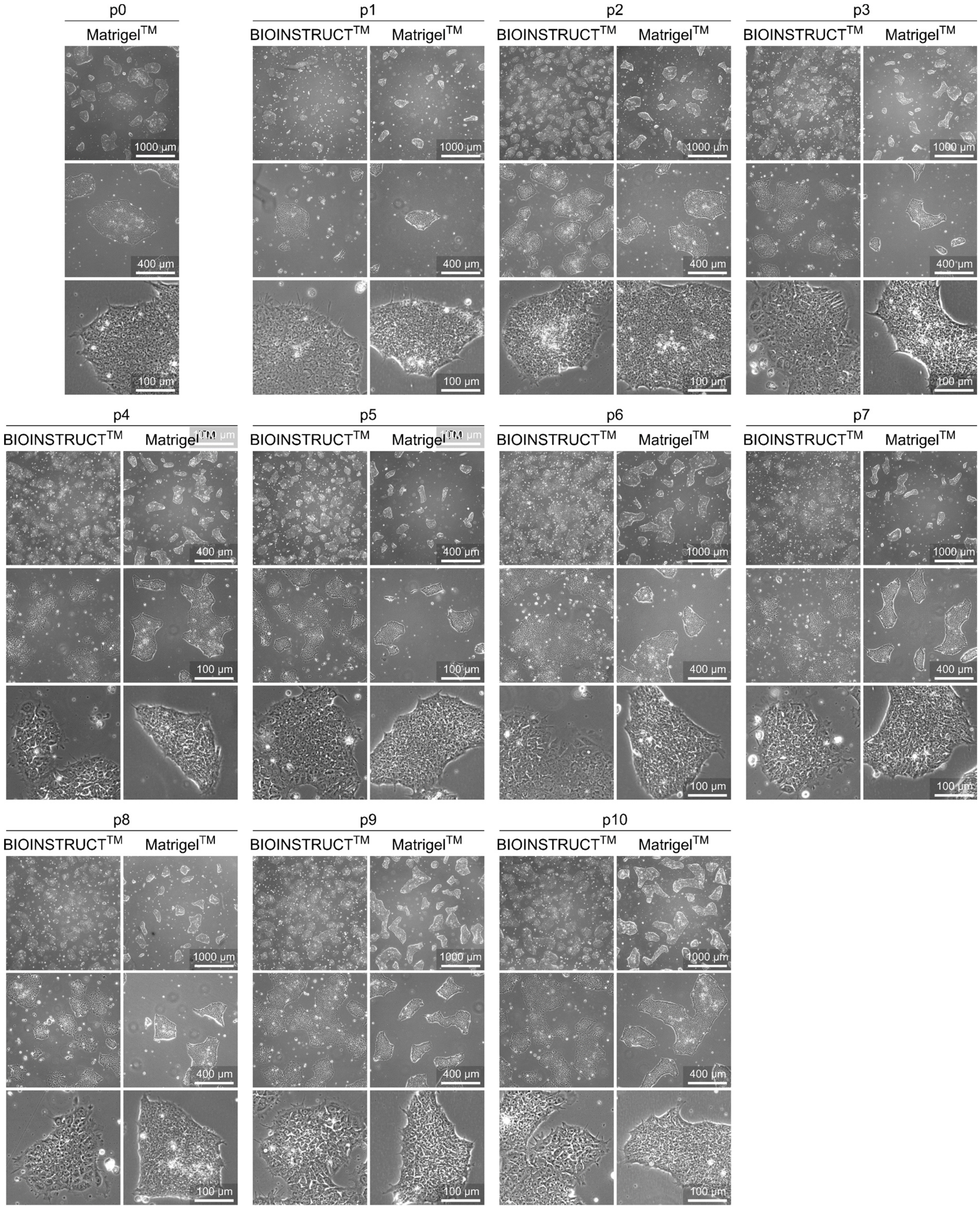


**Supplemental Figure 1 Cellular Morphology of hiPSC cultured on BIOINSTRUCT™ and Matrigel™ for ten passages.** Representative microscopic images of hiPSC cultured for ten on BIOINSTRUCT™ or Matrigel™. hiPSCs on both substrates form roundish, densely packed colonies and exhibit a small cell body with a high nucleus-to-cytoplasm ratio. Cells on BIOINSTRUCT™ appear slightly larger, with a slightly lower nucleus-to-cytoplasm ratio and form thin and flattened protrusions which are less frequently visible in cultures on Matrigel™.

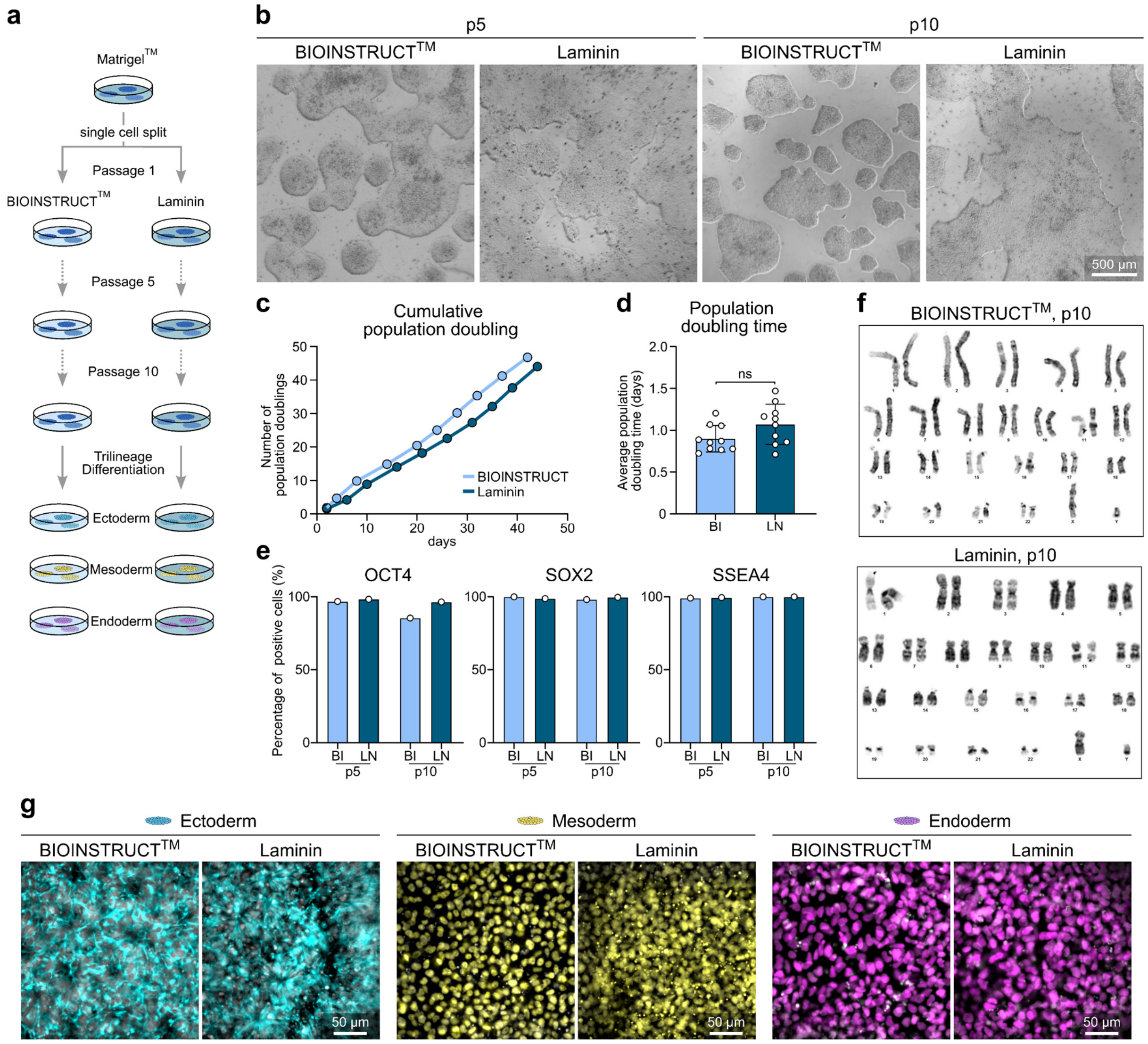


**Supplemental Figure 2 hiPSC cultured on BIOINSTRUCT™ retain a stem cell characteristic phenotype.** **a** Graphical illustration of the experimental set up. BIONi010B hiPSCs were cultured on BIOINSTRUCT™- or laminin-coated plates for 10 passages with single cell passaging every four to five days. **b** Brightfield images of hiPSCs cultured on BIOINSTRUCT™ or laminin at passage 5 and 10 after start of the experiment. **c** Cumulative population doubling of the hiPSC cultured on BIOINSTRUCT™ or laminin given as the number of population doublings over culture days. **d** Average population doubling time of hiPSCs cultured on BIOINSTRUCT™ or laminin. Data represent mean ± SD. Data points represent the population doubling time of single passages (n=10 from 1 experiment). Statistical analysis was performed using an unpaired two-tailed t-test. **e** FACS analysis of the core pluripotency genes *OCT4*, *SOX2*, and *SSEA4* of hiPSCs cultured on BIOINSTRUCT™ or laminin at passage 5 and 10. Data represent absolute percentage of positively stained cells (n=1 experiment). **f.** Karyogramm of hiPSC at passage 10 of culture BIOINSTRUCT™ or laminin. **g** Representative immunofluorescent images of trilineage differentiated hiPSCs cultured on BIOINSTRUCT™ or laminin at passage 10. Ectodermal cells are stained for Nestin (cyan), Mesodermal cells are stained for Brachyury (yellow), and Endodemal cells are stained for SOX17 (magenta). hiPSCs are counterstained with DAPI (grey). Abbreviations: BI: BIOINSTRUCT™, LN: Laminin, p: passage.

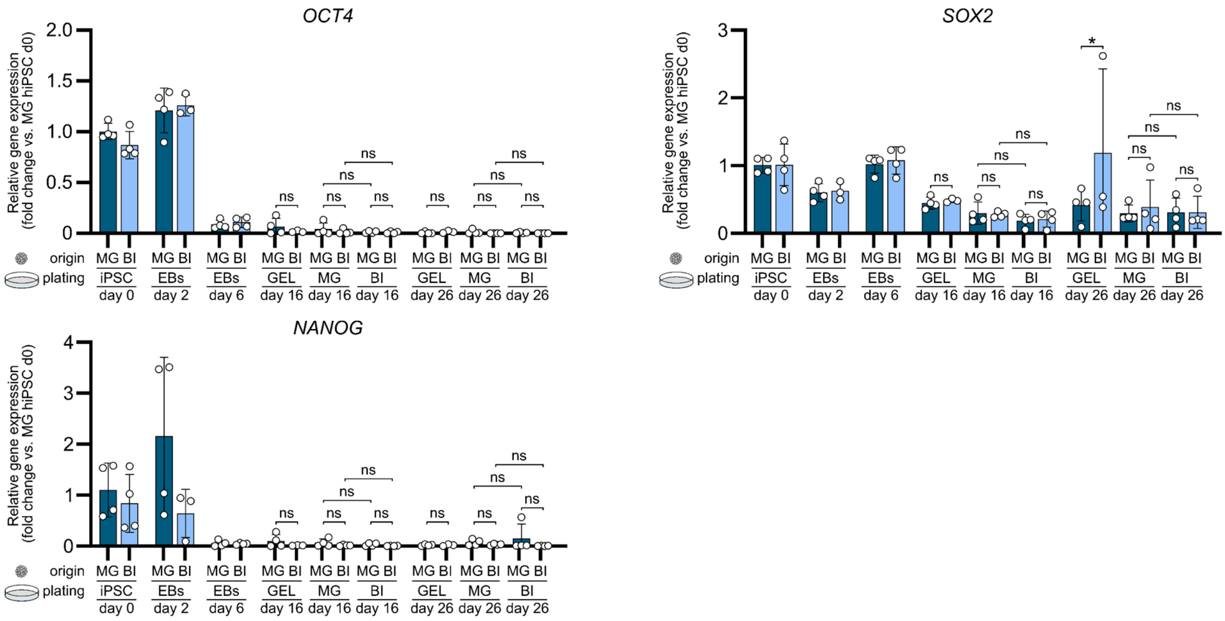


**Supplemental Figure 3 Gene expression analysis of pluripotency-associated genes during spontaneous differentiation on different cell culture substrates.** Bar graphs showing the expression of *OCT4*, *SOX2* and *NANOG* after eleven passages of BIOINSTRUCT$^{TM}$ or Matrigel$^{TM}$ (d0 hiPSC), after EB formation (EBs day 2), after 4 days exposure to spontaneous differentiation medium (EBs day 6) as well as ten (day 16) and 20 days (day 26) after plating on Gelatine, BIOINSTRUCT$^{TM}$ or Matrigel$^{TM}$. Gene expression is displayed as fold change to Matrigel d0 hiPSC. Data represent mean ± SD (n≥3 independent experiments). Statistical analysis was performed using one-way ANOVA followed by Sidak's multiple comparisons test. ns (not significant) P > 0.05.

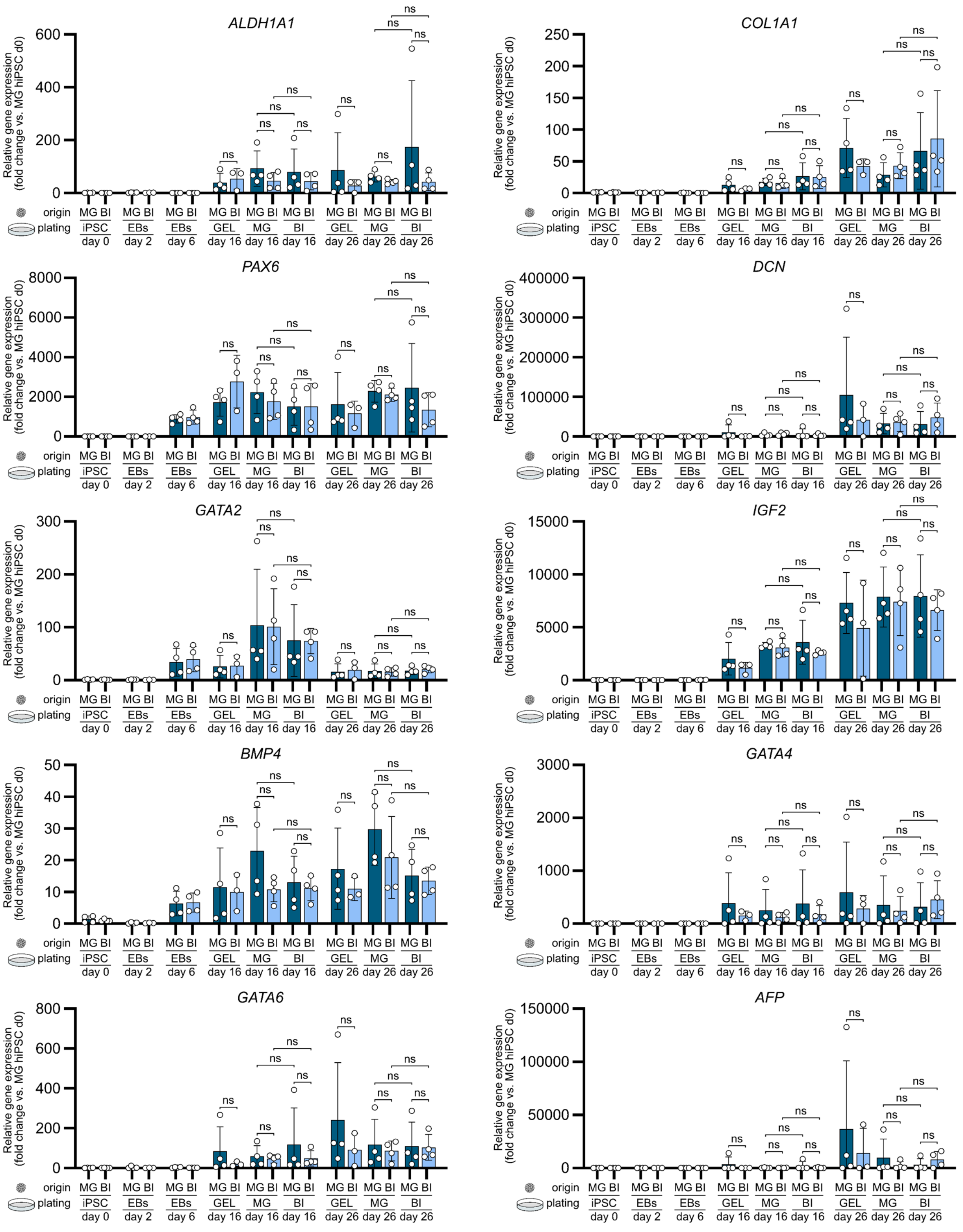


**Supplemental Figure 4 Gene expression analysis of genes associated with germ layer development in the course of spontaneous differentiation on different cell culture substrates.** Bar graphs showing the expression of the ectoderm-associated genes *ALDH1A1*, *COL1A1*, and *PAX6*, the mesoderm-associated genes *DCN*, *GATA2*, *IGF2*, and *BMP4*, and endoderm-associated markers *GATA4*, *GATA6*, and *AFP*. Marker assignment have been conducted according to the study by D'antonio et al.[24]. Gene expression is shown for cells after eleven passages of BIOINSTRUCT™ or Matrigel™ (d0 hiPSC), after EB formation (EBs day 2), after 4 days exposure to spontaneous differentiation medium (EBs day 6) as well as ten (day 16) and 20 days (day 26) after plating on Gelatine, BIOINSTRUCT™ or Matrigel™. Gene expression displayed as fold change to Matrigel d0 hiPSC. Data represent mean ± SD (n≥3 independent experiments). Statistical analysis was performed using one-way ANOVA followed by Sidak's multiple comparisons test. ns (not significant) $P > 0.05$, *$P < 0.05$.
