## Supplemental Document 2 for "A synthetic xeno-free extracellular matrix surrogate supports robust human induced pluripotent stem cell maintenance and multilineage differentiation"

### Document S2_Primer sequence and antibody information_prefinal

**Table 1** List of primers

| *Gene/RefSeq* | *Reference* | *T [C°]* | | *Primer sequence (5'-3')* |
|---|---|---|---|---|
| ***Housekeeper*** | | | | |
| *RPL4* NM_000968.4 | - | 60 | f | TATGCAAAGACCATGCGCC |
| | | | r | CCGGAGCTTGTGATTCCTGG |
| *RPL6* NM_000970.6 | - | 60 | f | GTGAGAAAACTGCGAGCCAG |
| | | | r | AGGAAAACCACCCTCTTGCC |
| *RPS29* NM_001032.5 | (D'Antonio et al. 2017) | 60 | f | AATATGTGCCGCCAGTGTTT |
| | | | r | CCCGGATAATCCTCTGAAGG |
| *GAPDH* | | 60 | f | GTCTCCTCTGACTTCAACAGCG |
| | | | r | ACCACCCTGTTGCTGTAGCCAA |
| ***Stem Cell characterization*** | | | | |
| *NANOG* NM_024865.4 | - | 60 | f | CCAAATTCTCCTGCCAGTGAC |
| | | | r | CACGTGGTTTCCAAACAAGAAA |
| *OCT4* NM_002701.6 | - | 60 | f | CCTCACTTCACTGCACTGTA |
| | | | r | CAGGTTTTCTTTCCCTAGCT |
| *SOX2* NM_003106.4 | - | 60 | f | CCCAGCAGACTTCACATGT |
| | | | r | CCTCCCATTTCCCTCGTTTT |
| ***Naive stem cell state*** | | | | |
| *KLF4* NM_001314052.2 | - | 60 | f | GCTGCCGAGGACCTTCTG |
| | | | r | GCGAACGTGGAGAAAGATGG |
| *DDPA3* NM_199286.4 | - | 60 | f | AGACCAACAAACAAGGAGCCT |
| | | | r | CCCATCCATTAGACACGCAGA |
| *KHDC3L* NM_001017361.3 | | 60 | f | AACCTGGCTGACTATCACCG |
| | | | r | GGGCCTTTCCTGAGCCTTTC |
| *HORMAD1* NM_032132.5 | | 60 | f | GCCCCTCAGACGTAATGTTG |
| | | | r | CACTCATGGGAGTCCTCTGC |
| *TRIM60* NM_152620.3 | | 60 | f | TGTGAGGGTTTCCGCTCTG |
| | | | r | GGGCAAACTGATGACCTGG |
| *DPPA5* NM_001025290.3 | | 60 | f | AGGTCGTGGTTTACGGCTC |
| | | | r | GCAAGTTTGAGCATCCCTCG |
| ALPG (ALPPL2) NM_031313.3 | | 60 | f | CTCTCCGACTGCTTCCAGAC |
| | | | r | GGTTCTCCTCCTCAACTGGG |
| ***Primed stem cell state*** | | | | |
| *ZDHHC22* NM_174976.2 | | 60 | f | CCGAGTCCACAACTTTGCAG |
| | | | r | CATCCTCGATTACATTCCGAGC |
| *ZIC2* NM_007129.5 | | 60 | f | GATGTGCGACAAGTCCTACAC |
| | | | r | TGGACGACTCATAGCCGGA |
| *THY1* NM_006288.5 | | 60 | f | CTCTCCTGCTAACAGTCTTGC |
| | | | r | TTAGGCTGGTCACCTTCTGC |
| *DUSP6* NM_001946.4 | | 60 | f | TTCCCTGAGGCCATTTCTTT |
| | | | r | AGTGACTGAGCGGCTAATG |
| *KLHL4* NM_019117.5 | | 60 | f | GTGATGCCTCCCATGTCAAC |
| | | | r | AGAGTGGCTACACCTAAGCC |
| *SOX11* NM_003108.4 | | 60 | f | CCTGTCGCTGGTGGATAAGG |
| | | | r | GTGCAGTAGTCGGGGAACTC |
| *PTPRZ1* NM_002851.3 | | 60 | f | TGTTTGCCGCCTGGATTGG |
| | | | r | TAGGACCAGCCAATCTCTTCAAC |

| | | | | |
|---|---|---|---|---|
| ***Germ Layer lineages*** | | | | |
| *AFP*<br>NM_001134.3 | (D'Antonio et al. 2017) | 60 | f<br>r | TGAGCACTGTTGCAGAGGAG<br>TTGTTTGACAGAGTGTCTTGTTGA |
| *ALDH1A1*<br>NM_000689.5 | (D'Antonio et al. 2017) | 60 | f<br>r | CTGCTGGCGACAATGGAGT<br>CGCAATGTTTTGATGCAGCCT |
| *BMP4*<br>NM_001202.6 | (Bock et al. 2011) | 60 | f<br>r | GCACTGGTCTTGAGTATCCTG<br>TGCTGAGGTTAAAGAGGAAACG |
| *COL1A1*<br>NM_000088.4 | (D'Antonio et al. 2017) | 60 | f<br>r | GAGGGCCAAGACGAAGACATC<br>CAGATCACGTCATCGCACAAC |
| *DCN*<br>NM_001920.5 | (D'Antonio et al. 2017) | 60 | f<br>r | AGAGTACCTGGTGGGCTGG<br>GTGGGCAGAAGTCACTTGAT |
| *GATA2*<br>NM_032638.5 | (D'Antonio et al. 2017) | 60 | f<br>r | CTGTCTGCAACGCCTGTG<br>GTTCCGAGTCTGGATCCCTT |
| *GATA4*<br>NM_001308093.3 | (Ruiz et al. 2010) | 60 | f<br>r | ACACCCCAATCTCGATATGTTTG<br>GTTGCACAGATAGTGACCCGT |
| *GATA6*<br>NM_005257.6 | (Ruiz et al. 2010) | 60 | f<br>r | AGGGCTCGGTGAGTCCAAT<br>CGCTGCTGGTGAATAAAAAGGA |
| *IGF2*<br>NM_000612.6 | (D'Antonio et al. 2017) | 60 | f<br>r | AGACGTACTGTGCTACCCC<br>TGCTTCCAGGTGTCATATTGG |
| *PAX6*<br>NM_001127612.3 | (Bock et al. 2011) | 60 | f<br>r | AACGATAACATACCAAGCGTGT<br>GGTCTGCCCGTTCAACATC |
| **Cardiac lineage markers** | | | | |
| *TTN*<br>NM_001267550.1 | (Yang et al. 2014) | 60 | f<br>r | GTAAAAAGAGCTGCCCCAGTGA<br>GCTAGGTGGCCCAGTGCTACT |
| *MYH6*<br>NM_002471.4 | (Baljinnyam et al. 2017) | 60 | f<br>r | TCAGCTGGAGGCCAAAGTAAAGGA<br>TTCTTGAGCTCTGAGCACTCGTCT |
| *MYH7*<br>NM_000257.4 | - | 60 | f<br>r | CCATTGGGGACCGCAGCAAGA<br>GCGGATTTGCCCAGGTGGTTG |
| *RYR2*<br>NM_001035.3 | - | 60 | f<br>r | TTGTTATGGCTCAGCTGTTTG<br>GCCTGGATGACAGTGTGAGA |
| *ATP2A2*<br>NM_001681.4 | (Johnson et al. 2021) | 60 | f<br>r | CGGAATTCATGCGCAGTGATAAATTGAC<br>CTCGGATCCAACACTACAGGTGTTGAATGG |
| **Neuronal lineage markers** | | | | |
| TUBB3<br>NM_006086 | | 60 | f<br>r | TGGACATCTCTTCAGGCCTG<br>ATGATGCGGTCGGGATACTC |
| MAPT<br>NM_005910 | | 60 | f<br>r | CCTCTGATCCAACCCTCCAG<br>TTTTGCTGGAATCCTGGTGG |
| MAP2<br>NM_002374 | | 60 | f<br>r | CCCAAGATACAGCTCAGCCT<br>GGAGGGAGAATGGAGGAAGG |
| BCL11B<br>NM_138576 | | 60 | f<br>r | GGTCTGGAGATAGAGGAGCC<br>TGTCCAGGGCCTTGTCATAG |
| TBR1<br>NM_006593 | | 60 | f<br>r | CCATCTGCCCATTGTTATTTG<br>CACCAATGTGCAAGGAAATC |
| SLC17A7<br>NM_020309 | | 60 | f<br>r | CGACGACAGCCTTTTGTGGT<br>GCCGTAGACGTAGAAAACAGAG |
| FOXG1<br>NM_005249 | | 60 | f<br>r | CAGACCTTGAGAACAACCTAAA<br>GGGCACCTTTACTACGAATG |
| SOX2<br>NM_003106 | | 60 | f<br>r | AGCTCGCAGACCTACATGAA<br>CCGGGGAGATACATGCTGAT |
| SYN1<br>NM_006950 | | 60 | f<br>r | CGATGCCAAATATGACGTGCGTG<br>AGCATCGCAGAGCCAGTATTGG |

| **Pancreatic lineage markers** | | | | |
|---|---|---|---|---|
| *ARX*<br>NM_139058.3 | | 60 | f | TCTCCTCCTACTGCATCGACA |
| | | | r | TCTTAGGGGAGCCTTGCAC |
| *GCG*<br>NM_002054.5 | (D'Amour et al. 2006) | 60 | f | AAGCATTTA CTTTGTGGCTGGATT |
| | | | r | TGATCTGGATTTCTCCTCTGTGTCT |
| *HHEX*<br>*NM_002729.5* | | | f | GGAGACTAAAACAGGAGAACCCT |
| | | | r | GGGAGGGCGAACATTGAGAG |
| *INS*<br>NM_000207.3 | | 60 | f | TCTACACACCCAAGACCCGC |
| | | | r | GTTCCACAATGCCACGCTTC |
| *MAFA*<br>NM_201589.4 | | 60 | f | AAAACCTTGGAGGAAGCCCC |
| | | | r | GAGCGACTCATGCTGTGACT |
| *NKX6.1*<br>NM_006168.3 | | 60 | f | TGGCCTGTACCCCTCATCAA |
| | | | r | CCGGAAAAAGTGGGTCTCGT |
| *PDX1*<br>NM_000209.4 | | 60 | f | CGTCCGCTTGTTCTCCTC |
| | | | r | CCTTTCCCATGGATGAAGTC |
| *SST*<br>NM_001048.4 | (Zhang et al. 2009) | 60 | f | CCCAGACTCCGTCAGTTTCT |
| | | | r | ATCATTCTCCGTCTGGTTGG |

**Table 2** List of primary antibodies

| ***Antigen*** | ***host*** | ***dil.*** | ***vendor*** | ***Catalogue #*** |
|---|---|---|---|---|
| CPEP | rt | 1:300 | DSHB | GN-ID4, supernatant |
| GATA2 | ms | 1:100 | Abcam | Ab22849 |
| GATA4 | rb | 1:250 | Abcam | ab5790 |
| GCG | ms | 1:1000 | Abcam | ab10988 |
| NANOG | rb | 1:150 | Abcam | ab80892 |
| NKX6.1 | ms | 1:100 | DSHB | F55A12, supernatant |
| OCT4 | gt | 1:100 | Abcam | ab27985 |
| PAX6 | rb | 1:250 | Abcam | ab124265 |
| PDX1 | gt | 1:250 | R&D | AF2419 |
| SOX2 | rb | 1:50 | Abcam | ab137385 |
| SST | rb | 1:500 | Sigma-Ald. | HPA019472 |
| TRA 1-60 | ms | 1:250 | Abcam | ab16288 |
| TUJ | ms | 1:1000 | Sigma | T8660 |
| CTIP2 | rat | 1:250 | Abcam | ab18465 |
| GFAP | rb | 1:500 | DAKO | Z0334 |
| Brachyury | ms | 1:200 | Invitrogen | MA5-14845 |
| Nestin | ms | 1:250 | BD Biosciences | 611659 |
| SOX17 | ms | 1:200 | BioLegend | 628152 |

**Table 3** List of secondary antibodies

| ***Antigen*** | ***Fluorophor*** | ***host*** | ***dilution*** | ***vendor*** | ***Catalogue #*** |
|---|---|---|---|---|---|
| gt | Alexa647 | dk | 1:400 | Life Technologies | A21447 |
| ms | Alexa555 | dk | 1:400 | Life Technologies | A31570 |
| rb | Alexa555 | dk | 1:400 | Life Technologies | A31572 |
| rt | Alexa488 | dk | 1:400 | Life Technologies | A21208 |
| ms | Alexa647 | gt | 1:1000 | Invitrogen | A11077 |
| rat | Alexa568 | gt | 1:1000 | Invitrogen | A21235 |
| rb | Alexa568 | gt | 1:1000 | Invitrogen | A11011 |

**Table 4** List of antibodies used for flow cytometry

| ***Antigen*** | ***Fluorophor*** | ***host*** | ***dil.*** | ***vendor*** | ***Catalogue #*** |
|---|---|---|---|---|---|
| Oct3/4 | PE | ms | 1:5 | BD Biosciences | 560186 |
| SSEA-4 | A647 | ms | 1:5 | BD Biosciences | 560796 |
| SOX2 | Cy5.5 | ms | 1:20 | BD Biosciences | 561506 |
| NANOG | APC | rb | 1:25 | Cell Signaling | 5448S |
| TRA1-81 | PE | ms | 1:25 | BD Biosciences | 560161 |
